# Iterative assay for transposase-accessible chromatin by sequencing to isolate functionally relevant neuronal subtypes

**DOI:** 10.1101/2023.04.14.536950

**Authors:** Collin B. Merrill, Iris Titos, Miguel A. Pabon, Austin B. Montgomery, Aylin R. Rodan, Adrian Rothenfluh

## Abstract

The *Drosophila* brain contains tens of thousands of distinct cell types. Thousands of different transgenic lines reproducibly target specific neuron subsets, yet most still express in several cell types. Furthermore, most lines were developed without *a priori* knowledge of where the transgenes would be expressed. To aid in the development of cell type-specific tools for neuronal identification and manipulation, we developed an iterative assay for transposase-accessible chromatin (ATAC) approach. Open chromatin regions (OCRs) enriched in neurons, compared to whole bodies, drove transgene expression preferentially in subsets of neurons. A second round of ATAC-seq from these specific neuron subsets revealed additional enriched OCR2s that further restricted transgene expression within the chosen neuron subset. This approach allows for continued refinement of transgene expression, and we used it to identify neurons relevant for sleep behavior. Furthermore, this approach is widely applicable to other cell types and to other organisms.

## INTRODUCTION

The brain is composed of many distinct types of neurons that form intricate connections across various regions. Each neuron type has specific properties, such as morphology, physiology, or gene expression, that influence local, circuit, and regional function.^1^ Thus, the ability to identify and target specific neurons is critical for understanding the networks and circuits that determine overall brain function. *Drosophila melanogaster* is an excellent model to study fundamental mechanisms in neurobiology, because vinegar fly brains consist of approximately 200,000 neurons and are less complex than mammalian brains. Still, even in this ‘simpler’ brain, single or bilateral pairs of neurons can affect behavior.^2, 3^ Further, neuronal cell type diversity is high, as indicated by the fact that ∼22,600 annotated neurons in the connectome fall into ∼5,600 distinct connectivity types.^4^ These findings underline the need for tools to identify, isolate, and manipulate neurons sparsely and specifically.

Previous large-scale efforts have been made to target groups of neurons in the *Drosophila* brain using stable transgenic lines.^5, 6^ Most of these methods have relied on shotgun approaches using random fragments or identified promoter regions from genes with known or predicted function in the nervous system fused to an exogenous transcription factor, Gal4.^6^ Using this approach, thousands of lines have been generated that have proven highly useful to the community. Still, drawbacks from this method include the limited ability to predict which neurons are targeted by this approach, thus necessitating the labor-intensive generation and screening of large numbers of transgenic lines. Furthermore, the collection of generated Gal4 lines includes many diverse neurons, but most lines are still expressed in hundreds of neurons across many distinct cell types.^6^ These findings raise the question of whether a data-driven approach that reveals the cell type-specific chromatin state might be useful to predictably generate tissue-specific tools.

Accessibility of neuronal chromatin is critical for proper gene expression.^7^ Previous single cell chromatin immunoprecipitation with sequencing (ChIP-seq) and Hi-C studies showed that chromatin accessibility is cell type-specific,^8–10^ even at the level of cell subtypes.^11, 12^ Assay for transposase-accessible chromatin by sequencing (ATAC-seq) is an approach that enables detailed and relatively straightforward genome-wide chromatin surveys.^13^ Indeed, ATAC-seq has been used to investigate how the chromatin landscape is involved in several cellular processes and diseases.^14–18^ Cells in the mouse hippocampus, for example, differ substantially in their chromatin accessibility, even within nominally similar cells such pyramidal cells, which form distinct clusters when analyzed by single cell ATAC-seq.^19^ It is less clear whether such differences in chromatin accessibility analyzed using ATAC-seq can be used to generate stable transgenic tools to target neuron subtypes selectively, but sparsely, in the brain.

We performed ATAC-seq to analyze the chromatin landscape between all tissues (whole adult flies) and neurons in the head (mostly the brain) selected by fluorescence-activated nuclear sorting. This analysis identified brain-enriched open chromatin regions (OCRs) that, when subcloned in front of a transgene, drove expression preferentially in the brain, though in distinct subsets of neurons. Conversely, whole body-enriched OCRs drove transgene expression largely outside the brain. Different brain-enriched OCR transgenes affected sleep behaviors when the OCR-expressing neurons were activated. To home in on the neurons responsible for a selected behavioral phenotype, we performed a second, iterative round of ATAC-seq specifically from round one transgene-expressing neurons that affected the selected phenotype. Subcloned round two OCR2s that were enriched compared to all neurons identified subsets of round one neurons that affected the selected behavioral phenotype when using intersectional genetics. Our results demonstrate that an iterative ATAC-seq approach can be used to isolate specific neuron subtypes that underlie distinct behavioral phenotypes in an informed manner. This data-driven approach thus provides an efficient technique to create genetic tools to identify and investigate the function of new neuronal populations that is translatable to other organisms and other tissues of interest.

## RESULTS

### Open chromatin regions drive tissue-specific gene expression

Gene expression is often used to define cell populations within complex tissues. In the brain, for example, inhibitory neurons are often identified by expression of marker genes for GABA signaling, such as *Gad1* or *vGAT*. Still, such markers are widely expressed in various cell subtypes, with active gene regulatory elements in enhancers providing spatial specificity to marker gene promoter activity.^20, 21^ Thus, we asked whether active OCRs outside of gene promoters would be able to drive gene expression. To answer this question and to ascertain whether OCR-driven gene expression could be done in a tissue-specific manner, we assayed OCRs in two very broad tissues—whole flies and neurons isolated from fly heads—using ATAC-seq.

We leveraged the *Gal4/UAS* system^22^ to drive nuclear GFP expression in whole flies using two independent whole-organism drivers (*Tubulin84B-* and *Actin5c-Gal4*; 4 samples per driver, 8 total whole-fly samples) and two neuron-specific drivers (*elav-* and *nsyb-Gal4*; 4 samples per driver, 8 total neuron samples). We isolated GFP-expressing nuclei from whole flies or from head neurons using fluorescence activated nuclei sorting, as described previously.^23^ The isolated nuclei were then used to generate ATAC-seq libraries. When we compared chromatin accessibility between whole flies and head neurons using unsupervised hierarchical clustering, the 8 samples grouped into two clear classes: head neurons and whole flies, with many more accessible OCRs observed in head neurons compared to whole flies (Fig. 1A).

**Figure 1.**
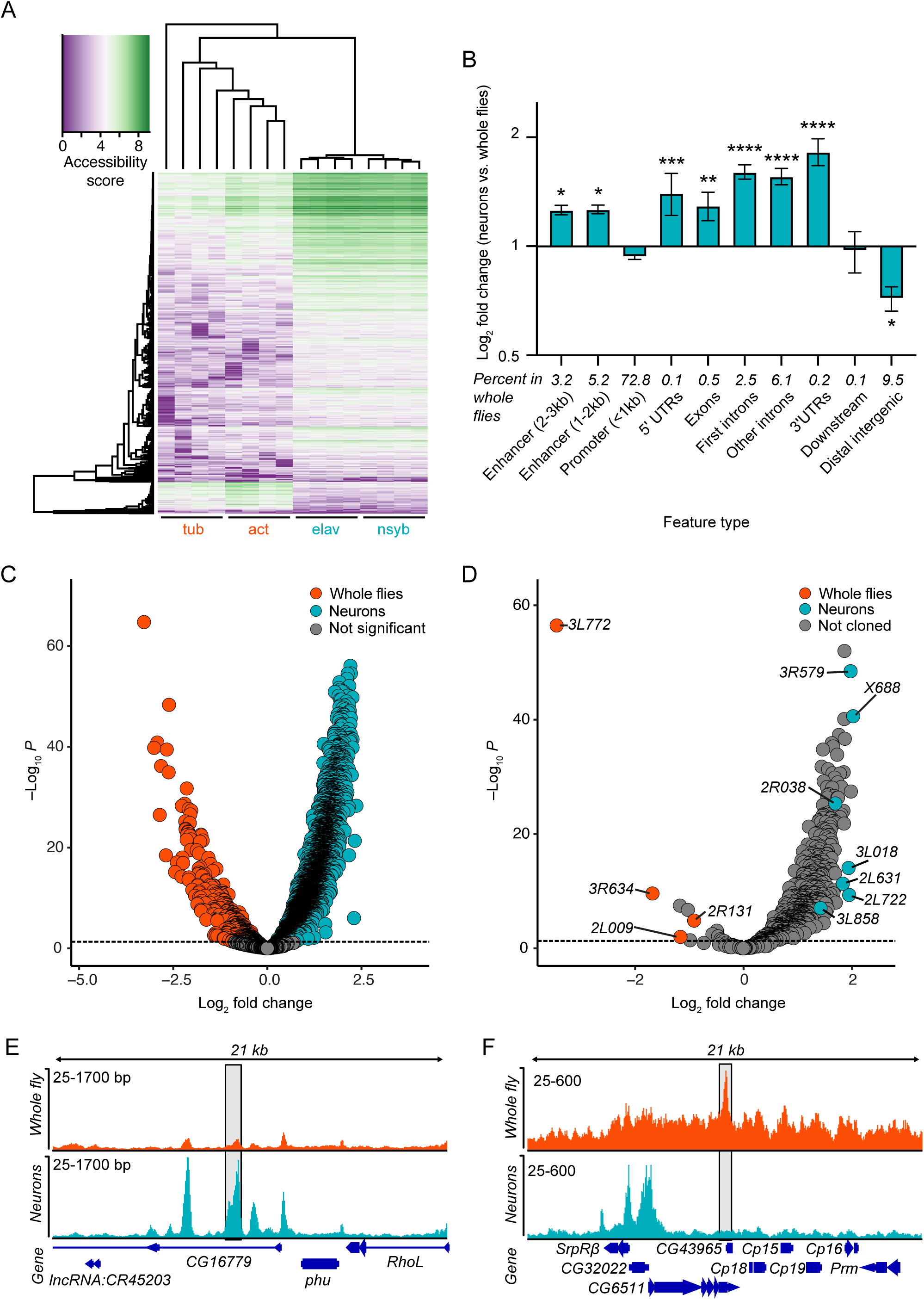
Differential chromatin accessibility between neurons and whole flies. A) Heatmap of chromatin accessibility in GFP-positive nuclei from whole flies (*Tubulin84B-* and *Actin5c-Gal4* drivers) and neurons from detached heads (*elav-* and *nsyb-Gal4* drivers). Each row represents a specific open chromatin region (OCR). Purple, decreased accessibility; green, increased accessibility. Each driver was analyzed using four biological replicates. B) Fold difference for fraction of OCRs annotated to each feature type in neurons compared to whole flies. The data were analyzed using one-way ANOVA followed by Holm-Sidak multiple comparisons tests. \**p* < 0.05, \*\**p* < 0.01, \*\*\**p* < 0.001 compared to whole flies. C) Volcano plot showing differentially-accessible OCRs from whole flies (orange) or neurons (cyan). The dotted line represents *p* = 0.05. D) Differentially accessible OCRs annotated to first intron or enhancer regions in whole flies (Log_2_ fold change < 0) and neurons (Log_2_ fold change > 0). The orange and cyan dots represent whole fly-and neuron-derived OCRs, respectively, that were used for subcloning experiments. E-F) Representative OCRs with increased accessibility in (E) neurons within gene *CG16779* and (F) whole flies within gene *CG43965*. The shaded area indicates the representative OCRs.

To determine whether specific gene regions or features were enriched in neuron-specific OCRs, we annotated each OCR (e.g. promoters, exons, introns, etc.) and calculated the proportion of OCRs annotated to each genomic feature type. We then compared the proportion of each feature in whole-fly and neuron OCRs. We found that the proportion of several feature types that typically contain regulatory sequences, including enhancers (2-3 kb from the transcription start site [TSS] and 1-2 kb from the TSS), 5′ untranslated regions (UTRs), exons, first introns, other introns, and 3′ UTRs were significantly higher in neurons compared to whole flies, while distal intergenic regions were lower (Fig. 1B). The increased proportions we observed for the enriched features in neurons suggests that these OCR types are particularly relevant for neuronal cell-type specific gene expression.

We next performed differential accessibility analysis on OCRs from whole flies and neuron samples. From the 16,337 consensus OCRs identified by pooling all libraries, we identified 12,988 differential OCRs. Of these, 439 were significantly more accessible in whole flies and 12,548 were more accessible in neurons (Fig. 1C). Because enhancers and first introns often contain regulatory elements in *Drosophila*,^24, 25^ we selected OCRs annotated to enhancers or first introns that were more accessible in whole flies (8 OCRs) or neurons (8 OCRs; 16 total OCRs; Fig. 1D-F). We subcloned the differentially-accessible OCRs upstream of a minimal *Drosophila* synthetic core promoter (*DSCP*)^5^ coupled to *Gal4* and injected the plasmids into fly embryos for insertion at the same genomic locus^26^ to generate stable transgenic *OCR*-*Gal4* fly lines. We named each line using the chromosome arm and last three digits of the sequence coordinate of the subcloned OCR (i.e. the peak located at <u>3R</u>:9490745-9491<u>579</u> was named *3R579*). Of the embryos injected with *OCR*-*DSCP*-*Gal4-*containing plasmids, we obtained 5 neuron OCR-derived lines (*2L722*, *3R579*, *X688*, *2R038*, and *3L858*) and 2 whole-fly OCR lines (*2R131* and *3R634*). Some injected *OCR-Gal4* plasmids (mostly whole fly-derived OCRs) did not yield stable transgenic lines, so we therefore repeated the OCR cloning approach using an *HSP70* minimal promoter instead.^27^ This yielded an additional 2 neuron-derived OCR lines (*3L018* and *2L631*) and 2 whole-fly OCR lines (*3L772* and *2L009*). All together, we generated 11 stable *OCR-Gal4* fly lines: 7 derived from regions with increased chromatin accessibility in neurons and 4 from whole flies.

### Differentially-accessible chromatin regions drive tissue-specific transgene activity

To test whether differentially-accessible OCRs drive tissue-specific *Gal4* expression, we first evaluated whole-body expression using a *UAS-GFP* reporter (Fig. 2A; Supplemental Fig. 1). Three *whole fly-OCR-Gal4* lines drove obvious-to-strong GFP expression in fly bodies, while *neuron-OCR-Gal4* lines all showed very dim GFP expression. Quantifying the GFP intensity (outside of the head) showed significantly higher expression in the bodies of *whole fly-OCR-Gal4* flies (*p =* 0.0006; *W* = 214, Wilcoxon signed-rank test; Fig. 2B). Similar results were obtained from whole-body sections stained with X-gal (from *OCR-Gal4;UAS-lacZ* flies; Supplemental Fig. 2). Quantifying the immunostaining intensity showed significantly more signal in brains from *neuron-OCR-Gal4* lines than from *whole fly-OCR-Gal4* lines (*p* = 0.003; *W* = 43, Wilcoxon; Fig. 2C; see Fig. 3 for immunostaining). Taken together, *whole fly-OCR* lines express more strongly throughout the body, while *neuron-OCR* lines express more strongly in brains (Fig. 2D).

**Figure 2.**
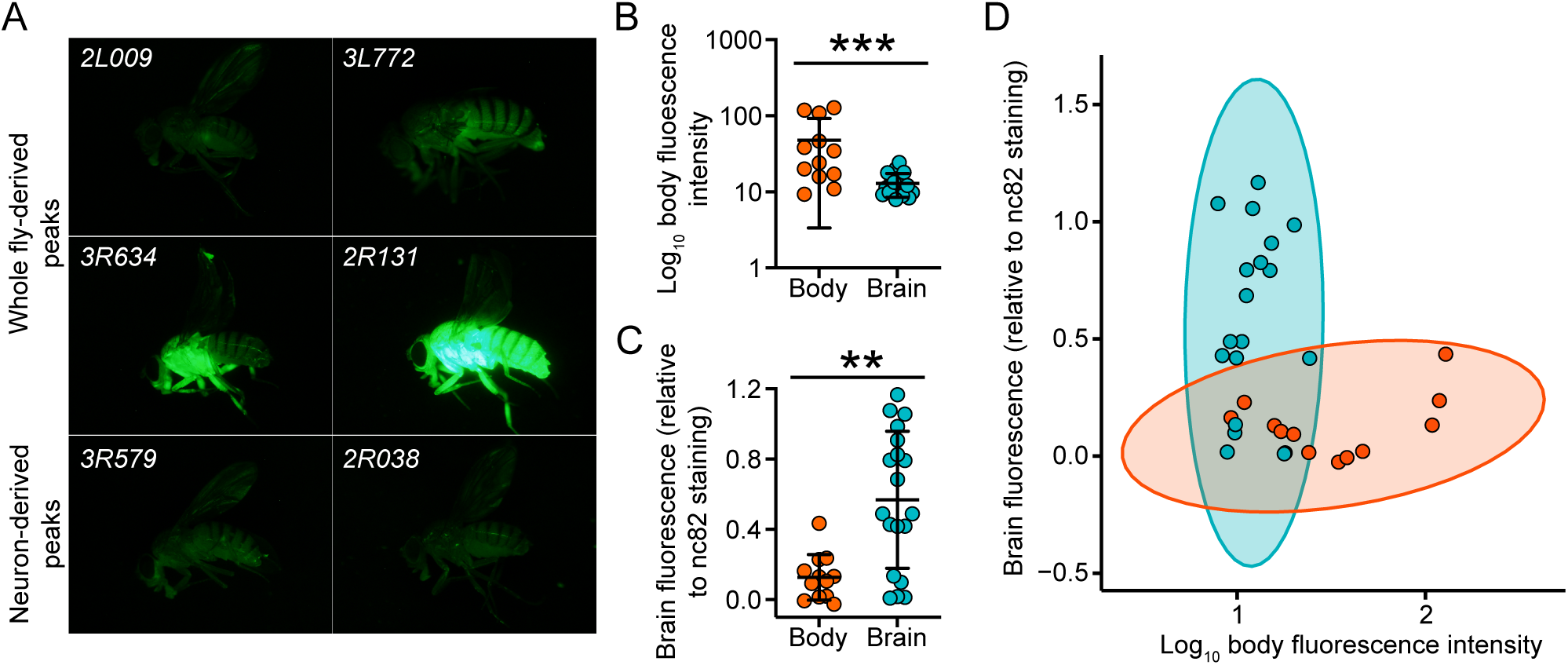
Chromatin regions that are more open in whole flies drive transgene expression in the fly body rather than the brain. A) Whole fly-derived OCR (*2L009*, *3L772*, *3R634*, and *2R131*) fragments drive Gal4 expression, which then activates *UAS-GFP*. GFP fluorescence is obvious in the body, while representative neuron-derived OCRs (*3R579* and *2R038*) drive little GFP in whole bodies. B) Quantification of GFP fluorescence in the body driven by whole fly-(orange) or neuron-derived (cyan) OCRs. C) Quantification of reporter expression in the brain driven by the distinct OCR-transgenes. D) GFP fluorescence driven by whole fly-vs. neuron-derived OCRs for each OCR. The data in B-C represent the mean ± SD and were compared using Wilcoxon signed-rank tests. \*\**p* = 0.003; \*\*\**p* = 0.0006. The fluorescence for immunostained reporters in the brain was normalized to nc82 reference staining.

**Figure 3.**
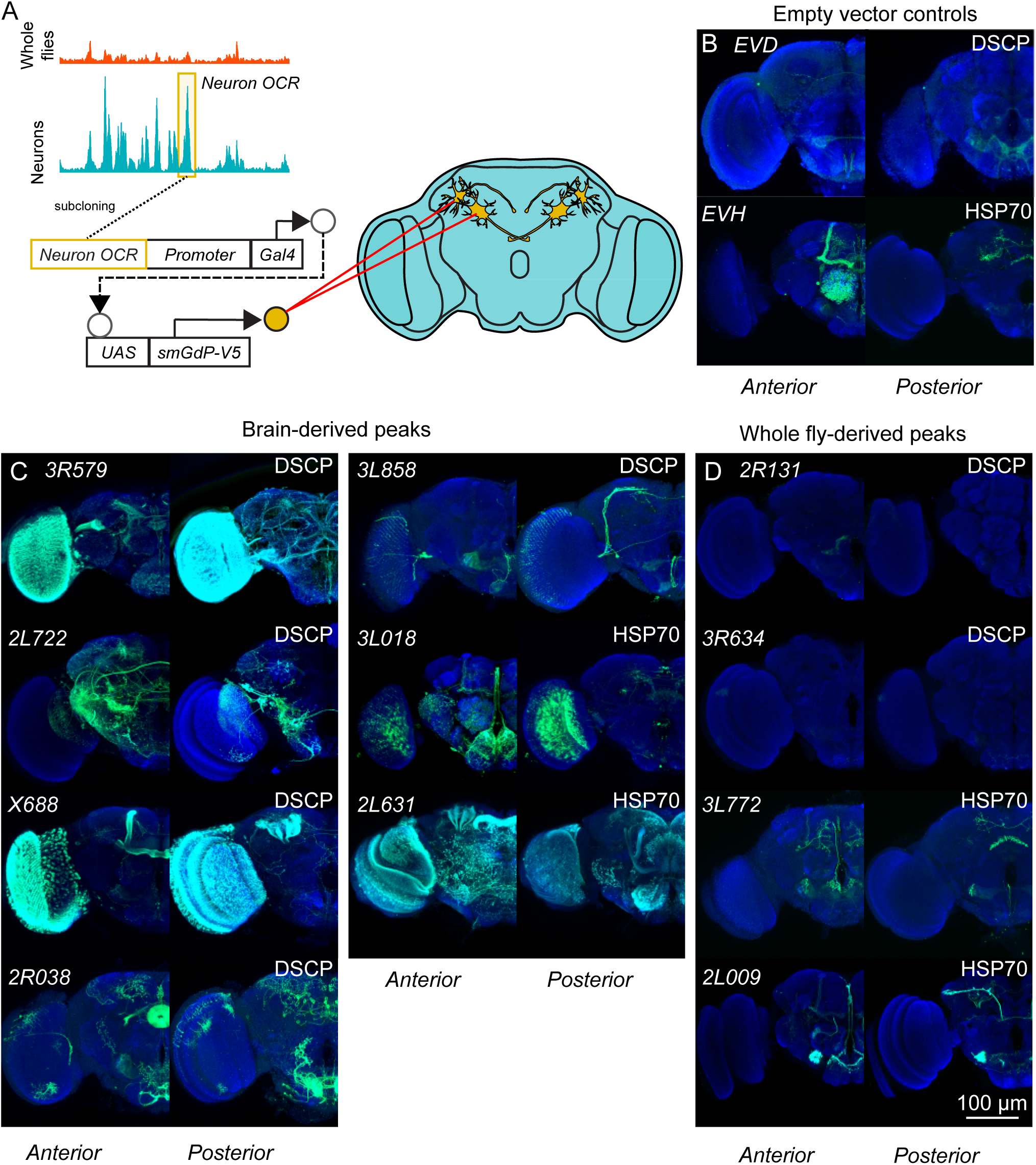
Neuron-derived OCRs drive transgene expression in different neurons. A) Schematic depicting ATAC-seq-derived *OCR* fragments 5′ to minimal promoters (*Drosophila* synthetic core promoter, *DSCP*, or promoter from the *Hsp70* gene) driving *Gal4*, which then activates membrane-bound myristoylated V5 (encoded by *UAS-myr::smGdP-V5*). B-C) Whole brain anti-V5 (green) and neuropil stainings (anti-nc82, blue) of B) empty vectors containing no OCR fragments, but just the promoters 5′ of *Gal4*, C) neuron-derived OCRs, and D) whole body OCRs. The OCR name is shown in the upper left corner of each image and the promoter is shown in the upper right corner. *EVD* and *EVH* in (B) indicate empty vectors containing the *Drosophila* synthetic core promoter or the minimal *HSP70* promoter, respectively. Scale bar indicates 100 µm, applicable to all images.

### Brain-specific open chromatin regions drive transgene expression in distinct neurons

Next, we examined transgene expression in the brain driven by *OCR-Gal4* by staining for a membrane-bound *UAS-GdP-V5* marker^28^ (Fig. 3A). Controls using the empty *DSCP* vector (*EVD*) or empty *HSP70* promoter (*EVH*) drove very sparse reporter expression (Fig. 3B), while each of the 7 *neuron-OCR-Gal4* lines drove V5 expression in several brain compartments, staining from thousands to tens of neurons (Fig. 3C). Conversely, *whole body-OCR-Gal4* lines drove markedly less V5 expression in the brain, with 2 of 4 lines showing no staining (*2R131* and *3R634*; Fig. 3D). The two *whole fly-OCR-Gal4* lines that showed expression in the brain (*3L772* and *2L009*) were both generated using vectors containing the *HSP70* minimal promoter. Interestingly, adding a *whole fly-OCR* recapitulated some empty vector staining (such as the fan-shaped body in *3L772*), drove marker expression in additional areas (such as antennal lobe projection neurons in *2L009*), but also subtracted some marker expression driven by the empty vector (i.e. antennal lobe intrinsic neurons that are absent in *3L772* and *2L009*), suggesting that the subcloned OCR regions contain instructive, as well as repressive, activity.

Aside from the striking difference in the number of neurons observed in *neuron-OCR-Gal4* lines, the staining patterns of these 7 lines were very diverse, with many labeled neurons expressed in distinct anatomical regions in one, but not other lines (e.g. the mushroom bodies in *X688*, the ellipsoid body in *2R038*, or the protocerebral bridge in *2L631*). These data show that neuron-derived accessible OCR regions drive expression in distinct subsets of neurons.

### Neurons identified by open chromatin regions drive behavioral phenotypes

The *Drosophila Gal4/UAS* system, used here to determine OCR-mediated reporter gene expression, is highly versatile and can be used to drive neuronal effectors, such as *UAS-TrpA1*, which allows for temperature-dependent neuronal activation at increased temperature (∼29 °C).^29^ Because we are interested in how individual neurons affect complex behaviors, we investigated how the *OCR-Gal4* neurons derived from our ATAC-seq analysis affected various sleep parameters. We chose this behavior because sleep is regulated by neural activity,^30, 31^ can be assessed for various phenotypes,^32^ and different neuron types are involved in sleep regulation.^33^

Out of 5 sleep parameters measured, only 1 of 4 *whole fly-OCR-Gal4* lines affected 1 of the 5 tested parameters (Fig. 4A), while 5 of 7 *neuron-OCR-Gal4* lines affected one or more parameters each (Fig. 4B). These data are consistent with our brain expression data, given that *neuron-OCR-Gal4* lines express more widely in the brain (Fig. 3), and show that neuron-derived accessible OCRs are more likely to be useful for identifying neurons that modulate behaviors.

**Figure 4.**
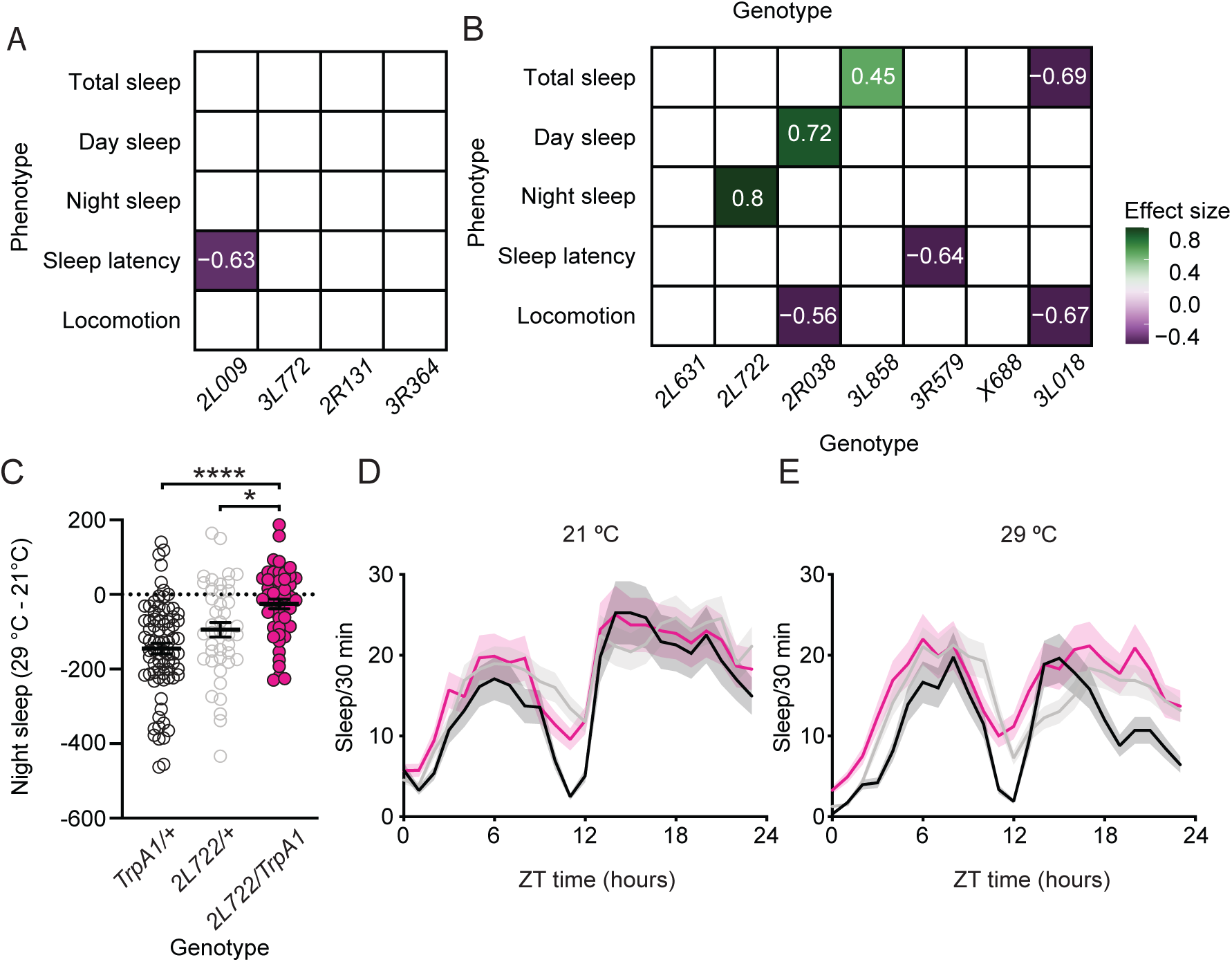
Neuron-derived OCRs express in neurons affecting diverse sleep phenotypes. Expression of temperature-activated TrpA1 channels (*UAS-TrpA1*) was driven using *OCR-Gal4*. Flies’ sleep measures for 2 days at 21 °C were compared to same flies for 2 days at 29 °C, which activates TrpA1 and the neurons expressing it. A,B) Effects on sleep phenotypes by activating TrpA1 are shown (Hedge’s *g* effect sizes are shown where significant, accounted for the effect of temperature in the absence of TrpA1 expression). (A) Whole fly-derived OCRs caused few effects on sleep phenotypes, while neuron-derived OCRs causes diverse sleep phenotypes (B). C) Temperature increases reduce night sleep duration in controls (left two genotypes), but not in *2L722>TrpA1* experimental flies, suggesting that *2L722*-expressing neurons promote night sleep (Kruskal-Wallis test followed by Dunn’s multiple comparisons test. \**p* = 0.01, \*\*\*\**p* < 0.0001). D-E) Average sleep-o-grams showing sleep at 21 °C (D) and 29 °C (E), with SEM shading.

### Iterative ATAC-seq identifies neuron-specific open chromatin regions that drive transgene expression

The *neuron-OCR-Gal4* line *2L722* is expressed in about 250 central brain neurons spanning many distinct brain regions (Fig. 3C). When these neurons were activated, flies showed increased nighttime sleep (Fig. 4C-E). Since the goal of our ATAC-seq approach was to develop tools for specific neuronal identification and manipulation, we asked whether we could further subdivide these 250 neurons in a data-driven manner and also identify smaller subpopulations that affect night sleep. *Neuron-OCR-Gal4s* were enriched for neuronal expression, but only targeted a subset of neurons. We thus reasoned that the same combination of specificity and sparsity might apply if we performed a second, iterative round of ATAC-seq, i.e. OCRs that were more accessible in *2L722* neurons would drive gene expression in a subset of *2L722* neurons.

To test this hypothesis, we performed ATAC-seq on GFP-labeled nuclei from two different *OCR-Gal4* lines: *2L722* and *3R579*. We detected 33,471 consensus peaks in pooled *2L722-*, *3R579-*, *elav-*, and *nsyb*-derived libraries. Principal component analysis (PCA) showed that the *2L722* and *3R579* samples clustered more closely to, but distinctly from, the neuronal samples than the whole-fly samples (Fig. 5A). Unsupervised hierarchical clustering showed that the overall chromatin accessibility was different between the whole fly, neuron, *2L722*, and *3R579* samples (Fig. 5B). Similar to the neuron vs. whole body comparison (Fig. 1B), *2L722* and *3R579* neurons showed a higher proportion of OCRs in enhancers, exons, introns, 5′ UTRs, and 3′ UTRs (Fig. 5C), suggesting that these regulatory regions confer increasingly cell-type specific gene expression.

**Figure 5.**
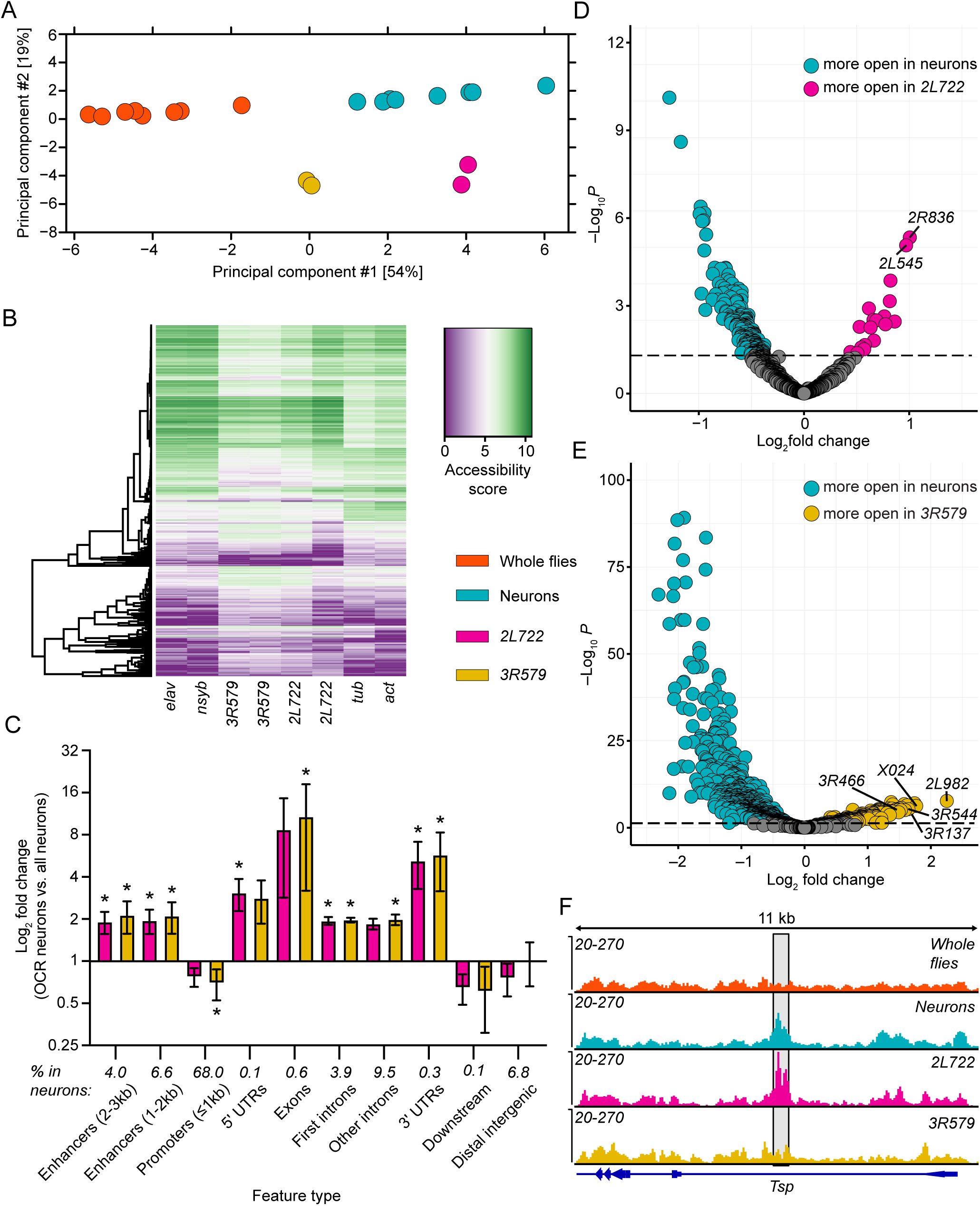
Iterative ATAC-seq identifies differentially-accessible chromatin regions in neuronal subtypes. A second round of iterative ATAC-seq was performed using GFP-positive nuclei from *2L722* and *3R579* neurons. A) Principal component analysis of OCRs from whole flies (orange), brains (cyan), *2L722* neurons (magenta), and *3R579* neurons (yellow). The x-and y-axes are scaled by the contribution of each principal component to the overall variability in the dataset. B) Heatmap showing the overall chromatin accessibility between whole flies, brains, *2L722* neurons, and *3R579* neurons. Purple indicates decreased accessibility and green indicates increased accessibility. C) Fold differences in the fraction of OCRs annotated to each genomic feature in *2L722* (magenta) and *3R579* neurons (yellow) compared to the proportion of each genomic feature in head neurons. The data were analyzed using Kruskal-Wallis tests followed by Dunn’s multiple comparisons tests, \**p* < 0.05 compared to head neurons. D) Volcano plot showing differentially-accessible OCRs in GFP-positive nuclei from neurons (cyan) and *2L722* neurons (magenta). D) Volcano plot showing differentially-accessible OCRs in GFP-positive nuclei from neurons (cyan) and *3R579* neurons (yellow). The annotated points in (D and E) represent subcloned OCR2s. F) A representative OCR annotated to the first intron of *Tsp* was identified in *2L722* neurons (magenta) and is more accessible than in *3R579* neurons (yellow), neurons (cyan), or in whole flies (orange). The shaded area indicates the OCR.

Differential accessibility analysis between *2L722* neurons and neuron-specific libraries (*elav* and *nsyb*) identified 6575 differentially accessible peaks (*p* < 0.05), with 487 peaks more open in *2L722* neurons and 6088 peaks more open in neurons. Comparing *3R579* neurons and whole brains identified 20,721 differentially accessible peaks (*p* < 0.05) in *3R579* libraries compared to neuron libraries, with 6283 more open regions in *3R579* neurons and 14,438 more open regions in neurons. Using the same approach we used to identify candidate OCRs in the first ATAC-seq experiment, we detected 1190 first intron OCR2s (for second round OCR) when comparing *2L722* neurons to whole brains, with 18 significantly more open in *2L722* libraries (Log_2_ fold change > 0; Fig. 5D). We also detected 1357 differentially-accessible first intron OCRs between *3R579* neurons and whole brains (Log_2_ fold change > 0). Of these, 183 peaks were more open in *3R579* neurons (Fig. 5E). We hypothesized that these second-round OCRs would identify a subset of the original neuron populations. Thus, we selected 2 and 5 significantly more accessible intron OCRs (OCR2) from *2L722* and *3R579* libraries (Log_2_ fold change > 1; Fig. 5F) and generated stable transgenic fly lines. OCR2s were subcloned upstream of the *DSCP* promoter to drive expression of *Flippase* (*FLP*) ^34^. We simultaneously used *2L722*-or *3R579-Gal4* and *OCR2-FLP* to express two reporter genes: 1) a *UAS*-*GFP* reporter driven by *2L722-* or *3R579-Gal4* in parental OCR neurons (Fig. 3A) and 2) an HA-tagged reporter downstream of *UAS* and the *FLP recombinase target (FRT)-STOP-FRT* sequence. Thus, the HA-tagged reporter was expressed only in neurons that simultaneously expressed *2L722-* or *3R579-Gal4* AND *OCR2-FLP*, since *2L722-* or *3R579-Gal4-*driven *FLP* expression causes the removal of the transcriptional *STOP* cassette in the *FRT-STOP-FRT*-*HA* reporter.

We observed sparse HA staining in the *OCR2-FLP*∩*2L722-Gal4* intersection (Fig. 6A-E). The staining was variable from fly to fly, likely due to incomplete penetrance of FLP-mediated excision of the *STOP* cassette, since FLP was originally intended to generate mosaic expression.^35, 36^ To increase FLP activity and resultant transgene expression, we heat-shocked the flies at the larval stage. This led to improved expression of the HA reporter, but expression was still clonal, i.e. often not bilaterally symmetrical and variable from individual fly to fly (of the same genotype). Despite this heterogeneity, we observed distinct intersectional *OCR2-FLP*∩*2L722-Gal4* staining between the *2R236-FLP* (Fig. 6B,C)*, 2L945-FLP* (Fig. 6D,E), and the control *empty-FLP* (*EVF*), which contained only a minimal *DSCP* promoter (Fig. 6F; Χ^2^ = 102.3, *p* = 5.19e-18, Chi-squared test; Supplemental Figure 4). *2R236-FLP*∩*2L722-Gal4* flies showed staining in the posterior dorsolateral protocerebrum projecting toward the protocerebral bridge (Fig. 6B,C), while *2L945-FLP*∩*2L722-Gal4* flies showed staining in the ventrolateral protocerebrum and the median bundle (Fig. 6D,E). In the *OCR2-FLP*∩*3R579-Gal4* lines, we observed sparse HA staining with some distinct intersectional staining in each *OCR2-FLP*∩*3R579-Gal4* line (Supplemental Fig. 5). Due to the complexity of the *3R579* expression pattern and the incomplete penetrance of FLP-mediated excision of the STOP cassette in the *OCR2-FLP*∩*3R579-Gal4* intersection, we were unable to reliably identify neurons stained for each intersection. However, the extant, but sparse staining we observed in the *OCR2-FLP*∩*3R579-Gal4* intersection supports our iterative ATAC-seq approach. Overall, these results show that – as hypothesized – our second round ATAC-seq-derived *OCR2-FLP* lines express transgenes in subsets of the parental OCR neurons and that different *OCR2-FLP* lines are comprised of distinct subsets of OCR neurons.

**Figure 6.**
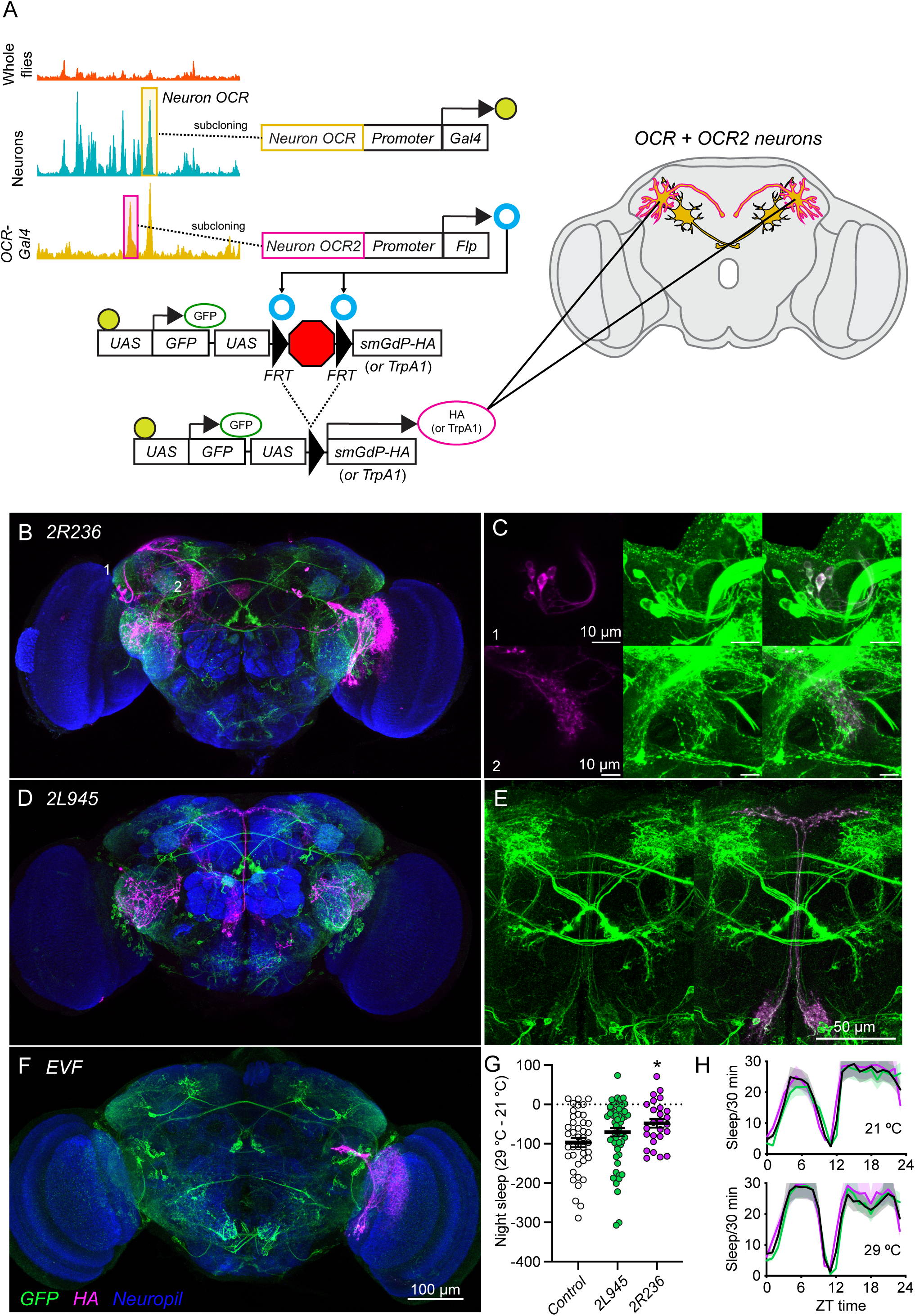
Differentially-accessible OCRs detected by iterative ATAC-seq drive transgene expression in *2L722* neuron subtypes. A) Schematic depicting the subcloning and genetic strategy. B-F) GFP expression (green) driven by first round *2L722-Gal4* and HA expression (magenta) driven in neurons expressing both *2L722-Gal4* and *2L722*-derived iterative *OCR2-FLPs*. Neuropil (nc82) is stained in blue. B) *2L722*∩*2R236* neurons. C) Zoomed images showing the cell bodies (1) and projections (2) of HA-positive lateral horn neurons isolated with *2L722*∩*2R236* intersectional drivers. D) *2L722*∩*2L945* neurons. E) Zoomed images showing *2L722*-driven GFP only (left) and *2L722*∩*2L945-*driven HA colocalization in median bundle neurons as well (right). F) *2L722*∩*Empty vector FLP* (*EVF*, with *DSCP* promoter), which stains in some optic lobe neurons. G) Temperature increases reduce night sleep duration in controls (black) and *2L722*∩*2L945* activated neurons (green), but significantly less so in *2L722*∩*2R236* neurons (blue), suggesting these intersectional neurons (B) promote night sleep. \**p* = 0.04, Kruskal Wallis test followed by Dunn’s multiple comparisons test. G) Average sleep-o-grams showing nighttime sleep at 21 °C and 29 °C.

**Figure 7.** Iterative ATAC-seq isolates behaviorally-relevant neuron subtypes. ATAC-seq 1 identifies neuron-specific OCRs comparing broad tissue types (i.e. whole flies vs. head neurons). Neuron-specific OCRs subcloned upstream of *Gal4* (or other reporter genes) drive transgene expression in distinct neuron subpopulations. Analyzing OCRs within these neuronal subpopulations by iterative ATAC-seq (round 2; black arrow) identifies OCRs that are overrepresented in that subpopulation. These OCR2s can be subcloned upstream of *FLP* (or other combinatorial gene expression systems) to home in on a progressively smaller neuronal subpopulation. This iterative ATAC-seq is translatable to other tissues or organisms.

### Chromatin regions with increased accessibility identified by iterative ATAC-seq impart subtype-specific behavioral phenotypes

Finally, we examined whether the *OCR2-FLP*∩*2L722-Gal4* intersectional neurons have functional relevance. To do so, we crossed the *2R236-FLP*∩*2L722-Gal4* and *2L945-FLP*∩*2L722-Gal4* lines to *UAS-FRT-STOP-FRT-TrpA1* so that only *FLP*-AND *Gal4*-positive neurons expressed TrpA1, which allowed us to activate *OCR2*-containing intersectional neurons in a temperature-dependent manner. Our hypothesis was that a subset of *2L722* neurons is sufficient to mediate the TrpA1-induced increase in nighttime sleep (Fig. 4). While *2R945-FLP*∩*2L722-Gal4* flies had the same night sleep as the *empty-FLP*∩*2L722-Gal4* controls, *2R236-FLP*∩*2L722-Gal4* flies showed an increased night sleep phenotype (F(2,120) = 3.566, *p =* 0.03; \**p* = 0.02, Dunnett’s multiple comparisons test; Fig. 6G,H), which was similar to the increased nighttime sleep phenotype observed in the original *2L722-Gal4;UAS-TrpA1* flies. These results indicate that *2R236-FLP*∩*2L722-Gal4* neurons, a subset of the original ∼250 *2L722* neurons, are involved in promoting night sleep. Thus, our approach of starting with a given tissue (neurons) and performing ATAC-seq to generate specific, but sparse, transgenic lines can be successfully used iteratively to isolate functionally relevant, sparse neuron subpopulations. Further, our approach can be used to isolate many other behaviorally relevant neuronal subpopulations in other tissues or in other organisms.

## DISCUSSION

The brain consists of thousands of different cell types, with many yet to be investigated regarding their function. Because of this vast diversity, we need tools – including new approaches for the efficient generation of such tools – to identify and study many of these neuronal cell types. Here we show that tissue-specific, iterative ATAC-seq is a useful approach to generate such new tools.

We determined genome-wide OCRs for the whole body and for the brain. Many more OCRs were identified in the brain compared to the whole body. When we determined genome-wide OCRs from specific neurons, we again identified more OCRs in specific neurons compared to all neurons. This is likely due to the ATAC-seq signal-to-noise ratio, since an OCR that is accessible in a specific subset of neurons may not result in a large enough signal to be detected when surveying all neurons, much less all cells in the body. Similar results were found when comparing mouse bulk cortex to more specific entorhinal cortex data, albeit with data derived from histone mark-specific chromatin immunoprecipitation, which resulted in more enhancers being identified in the more specific tissue.^37^ In our data, the OCRs from more specific tissues encompassed a larger proportion of OCRs in enhancers and throughout the transcriptional unit, while a larger proportion of accessible OCRs were located in promoters in more general tissue samples. This suggests that non-promoter regions confer tissue-specific gene expression and could be used to generate tools to identify and manipulate subtypes of cells. Indeed, enhancer regions are cell-specific ^38^ and are critically involved in regulating promoter activity and downstream gene expression.^39^

When we subcloned first intron OCRs that were more accessible in neurons compared to whole flies, we found that they drove transgene and reporter expression preferentially in neurons. Conversely, OCRs that were enriched in whole fly-derived data drove very sparse or no expression in the brain. Thus, OCRs that are enriched in a specific tissue drive expression preferentially in the tissue of interest. This is consistent with mammalian studies showing that tissue-specific ATAC-seq can be used to generate tissue-specific tools. These approaches have yielded viral AAV vectors that express in subsets of GABAergic interneurons starting from GABA neuron-specific ATAC-seq ^40^ or in subsets of the visual cortex starting from adult primary visual cortex single cell ATAC-seq.^41^ Therefore, tissue specific ATAC-seq data can be harnessed to generate tissue-specific tools in *Drosophila* and mammals.

Since the neuron-specific transgenes we generated are expressed in tens to thousands of neurons, we re-applied the same approach and performed a second, iterative ATAC-seq experiment to determine a second round of OCRs with greater accessibility in a subset of neurons. We then subcloned these second round OCRs so we could use *Gal4/FLP* intersectional genetics. This approach yielded new tools that were expressed in a subset of a set of neurons, showing that this iterative approach can be used to isolate increasingly specific cell populations. Compared to our first round of ATAC-seq from whole bodies and neurons (Fig. 1), we obtained many fewer differentially accessible OCRs when we compared pan-neuronal vs. specific neuron ATAC-seq (Fig. 5). Thus, we only generated 2 second round *OCR2-FLP-transgene* lines to dissect the parental *2L722-Gal4* pattern. To more systematically dissect the 2L722 pattern, one could generate more second round transgenic lines and increase sequencing depth to identify additional differentially accessible OCRs to begin with.

Furthermore, our FLP approach yielded more clonal intersectional patterns than hoped for. We used FLP because that approach is compatible with intersectional genetics using any parental Gal4 line. Alternatively, the split Gal4 system could be used for more reproducible intersectional patterns,^42^ but this approach would require *parent-Gal4* lines to be converted into *parent-split-Gal4* lines, which is not always straightforward. In general, however, an intersectional genetic approach to further subdivide a pre-existing expression pattern is also available in mammals using recombinases such as Cre or even FLP. Indeed, such an approach has been used in the mouse brain to co-inject and differentially/combinatorially label two distinct ATAC-seq-derived viruses.^41^ Our novel iterative ATAC-seq approach can thus be used to sequentially home in on decreasing subsets in a tissue of interest, and one could even perform a third iterative round of ATAC-seq data acquisition and OCR3 subcloning for additional specificity. Furthermore, tissue-specific ATAC-seq and intersectional strategies for in vivo expression are available from flies to mammals.

We also show that our approach can be used to isolate functionally relevant neurons. From our first round *OCR-Gal4* transgenes, more of the neuron-enriched lines affected sleep behavior then from the body-enriched ones (Fig. 4), as one might predict, given that the *neuron-OCR-Gal4* lines are more widely expressed in the brain than the *body-OCR-Gal4* lines (Fig. 2,3). Our intersectional approach (Fig. 6) suggested that neurons in the lateral horn (Fig. 6B,C) are involved in the nighttime sleep phenotype we observed in the parental *2L722-Gal4* line (Fig. 4). While we have not confirmed the identity of these putative sleep-regulating neurons, it is worth noting that some lateral horn neurons are involved in the regulation of sleep ^43^ and output from the circadian clock ^44^. Thus, our approach can be leveraged to identify and manipulate increasingly specific sets of neurons that are behaviorally relevant. The iterative nature of our approach represents a key step forward in the generation of increasingly cell type-specific tools and can be applied to any tissue of interest across the phylogenetic tree.

## ACKNOWLEDGMENTS

We thank members of the Rothenfluh and Rodan labs for discussion. We acknowledge the Cell Imaging Core at the University of Utah for use of equipment (Leica SP8 White Light laser confocal microscope). Stocks obtained from the Bloomington *Drosophila* Stock Center (NIH P40OD018537) were used in this study. This work was supported by the University of Utah Flow Cytometry Facility, the University of Utah Genomics Core Facility, the High Throughput Sequencing Core at the Huntsman Cancer Institute, and the National Cancer Institute through Award Number 5P30CA042014. This study was supported by grants from the National Institute of Health: National Institute on Drug Abuse (Grant R21DA049635, to AR), the National Institute on Alcohol Abuse and Alcoholism (K01AA029200 to CBM; R01AA026818 and R01AA019526 to AR), and the National Institute of Diabetes and Digestive and Kidney Disease (R01DK110358 to ARR). The content is solely the responsibility of the authors and does not necessarily represent the official views of the National Institutes of Health.

## AUTHOR CONTRIBUTIONS

Conceptualization: C.B.M and A.R.; Formal analysis: C.B.M, I.T., A.B.M., and A.R.; Investigation: C.B.M., I.T., and M.A.P.; Resources: M.A.P.; Data curation: C.B.M. and A.B.M.; Writing—Original draft: C.B.M. and I.T.; Writing—Review and editing: C.B.M., I.T., M.A.P., A.R.R., and A.R.; Visualization: C.B.M.; Supervision: A.R.; Funding acquisition: C.B.M., A.R.R., and A.R.

## DATA AND REAGENT AVAILABILITY

All sequencing data are deposited in the Gene Expression Omnibus database (Accession number: GSE226514). All generated plasmids and fly lines are available upon request.

## DECLARATION OF INTERESTS

The authors declare no competing interests.

## STAR METHODS

### Fly strains

The following fly lines were obtained from the Bloomington Drosophila Stock Center (Indiana University, Bloomington, IN, USA) and used in this study: *Actin5c-Gal4* (BL25374), *Tubulin84B-Gal4* (BL5138), *elav*-*Gal4* (BL458), *nSyb-Gal4* (BL51635), *UAS-GFP-nls* (nuclear localization signal; BL4775), *UAS-Stinger* (a super-bright GFP-nls variant; BL84277), *UAS-mCD8::GFP* (BL32184), *UAS-myr::smGdP-V5* (BL62146), *UAS-LacZ* (BL1776), *UAS-TrpA1* (BL26263), *UAS-myrGFP UAS-FRT-STOP-FRT-smHA* (BL62128), and *UAS-FRT-STOP-FRT-TrpA1* (BL66871). Flies were reared in bottles containing standard cornmeal agar and grown at 25 °C with 70% relative humidity and a 12-h light/dark cycle.

### ATAC-seq library construction

Flies containing *Tubulin84B-Gal4*, *Actin5c-Gal4*, *elav-Gal4*, or *nsyb-Gal4* drivers were used to induce nuclear GFP expression in whole flies or neurons. GFP-positive nuclei were isolated using our previously-described workflow.^23^ The isolated nuclei were resuspended in 1 mL wash buffer (10 mM Tris-HCl, 10 mM NaCl, and 3 mM MgCl_2_) containing 3 μM DAPI and were sorted using a BD FACS Aria III flow cytometer (BD Biosciences, San Jose, CA, USA) operated by the University of Utah Flow Cytometry Core facility. GFP-negative nuclei collected from *w* Berlin* flies (no GFP) were stained with 3 μM DAPI and used as the GFP-negative control to set the sorting gates. Nuclei were identified based on the forward scatter signal and DAPI intensity. GFP-positive nuclei were collected into 500 μL wash buffer and stored on ice until use for ATAC-seq library prep. The purified nuclei were used for library preparation as previously described,^23, 45^ with tagmentation for 23 min at 37 °C using 1X Tn5 enzyme (Illumina, Inc., San Diego, CA, USA). Fragments were amplified with Nextera index primers (Illumina) and Phusion HiFi master mix (New England Biolabs, Ipswich, MA, USA) for 5 cycles, followed by another 9-10 cycles per sample (as determined from 25% of total fluorescence from a qPCR side reaction performed using SsoFast EvaGreen SuperMix, BioRad, Hercules, CA, USA). The DNA fragments were purified with 0.5X and 1.3X AMPure XP beads (Beckman Coulter, Indianapolis, IN, USA) to remove primer-dimers and high-molecular weight DNA (> 5000 bp). The purified DNA was stored at −20 °C until further analysis.

### Sequencing and analysis

ATAC-seq libraries constructed using nuclei from whole flies (*Tubulin84B-Gal4* and *Actin5c-Gal4*) and head neurons (*elav-Gal4* and *nsyb-Gal4*) were sequenced on a HiSeq 2500 instrument using 50-bp paired-end reads. The fastq files were quality checked using FastQC (v 0.11.9). Adapter sequences were removed with CutAdapt (v3.4).^46^ The sequenced reads were aligned to the dmel_r6.26 genome assembly using Bowtie2 (v2.4.2).^47^ Aligned reads were sorted with Samtools (v1.12)^48^ and deduplicated with Picard (v2.23) using the MarkDuplicates command. Aligned, filtered, and deduplicated reads were used to call peaks using MACS2 software (v2.2.5).^49^ Differential accessibility analysis was performed using DiffBind (v2.10.0)^50^ and DESeq2 (v1.32.0).^51^ Peak annotation was performed using ChIPseeker (v1.30.3).^52^ Coverage files were generated using DeepTools (v3.3.2).^53^ Peaks were visualized using the IGV browser (v2.7.2).^54^

### Plasmid construction

Unless otherwise stated, all restriction enzymes were from NEB. All PCR amplification steps were performed using Phusion High Fidelity PCR MasterMix (NEB) and the primers listed in Supplemental Table 1 (Invitrogen). Amplified fragments were purified from 1% agarose gels using Monarch DNA Gel Extraction Kits (NEB). A multiple cloning site (MCS; Supplemental Table 1) containing unique PacI, NheI, StuI, and AvrII restriction sites was designed and synthesized as a gBlock (Integrated DNA Technologies, Coralville, IA, USA). The MCS was subcloned into pENTR_D_TOPO (ThermoFisher) and was inserted into pBPGw (17574, Addgene) using LR Clonase II (ThermoFisher). The resulting plasmid (pBP-MCS-DSCP-Gal4; pMDG) was digested with AvrII and re-ligated with T4 DNA ligase (NEB) to remove a 90-bp filler sequence within the MCS. To build pBP-MCS-Hsp70-Gal4 (pMHG), the 292-bp *Hsp70* promoter was amplified from pT-Arf6-T44N-GFP (an in-house plasmid) with the primers listed in Supplementary Table 1. The *HSP70* promoter was amplified, gel-purified, and subcloned into FseI/KpnI-digested purified pMDG using NEBuilder HiFi DNA assembly mix (NEB). pBP-MCS-DSCP-FlpD5 plasmid (pMDF) was constructed by amplifying the FlpD5 sequence from pBPhsFlp1 (32148, Addgene) with the primers listed in Supplementary Table 1. In parallel, pMDG was amplified with primers (Supplementary Table 1) designed to remove the Gal4 coding sequence. This yielded a linear backbone for FlpD5 insertion. The linear backbone and FlpD5 were assembled with NEBuilder HiFi DNA Assembly Mix.

To construct plasmids containing *OCR*s, genomic DNA was purified from whole *w* Berlin* flies using Monarch Genomic DNA purification kits (NEB). *OCR*s with significantly increased accessibility in neurons and whole flies were selected using the following criteria: largest log_2_ fold change, distance > 500 bp from the TSS, and *OCR* location within the first intron or within a promoter 2-3 kb from the TSS. *OCR* sequences plus 200-bp flanking regions were amplified from genomic DNA with primers that included 20-bp sequences complementary to the sequences upstream from the PacI and downstream from the AvrII sites in pMDG, pMHG, and pMDF (primer sequences in Supplemental Table 1). Amplified DNA fragments were gel purified and subcloned into PacI/AvrII-digested and gel-purifed pMDG, pMHG, and pMDF using NEBuilder HiFi DNA assembly mix (NEB). The assembled plasmids were Sanger sequenced to confirm correct assembly (Genewiz).

### Fly injection

The *OCR* plasmids were injected into P{y[+t7.7]=nos-phiC31\int.NLS}X, y[1] sc[1] v[1] sev[21]; P{y[+t7.7]=CaryP}attP2 flies (BL25710). The *OCR2* plasmids were injected into y[1] v[1] P{y[+t7.7]=nos-phiC31\int.NLS}X; P{y[+t7.7]=CaryP}attP40 flies (BL25709). All injections were performed by Rainbow Transgenic Flies, Inc.

### OCR-Gal4 characterization

*OCR-Gal4* flies were crossed to *UAS-Stinger* virgins. Adult progeny were examined for GFP expression using an Olympus SZX10 dissecting microscope equipped with an Olympus DP72 camera. Images were captured using Olympus CellSens software (RGB color mode, exposure time = 300 ms, and ISO sensitivity = 800). For each genotype, images from three different females were collected. GFP fluorescence in whole flies (excluding the head) was determined using ImageJ software. To corroborate the ImageJ analysis, each image was printed in duplicate and assigned a random number, which resulted in six numbered images per genotype. One image per genotype was used to generate 6 image decks (11 images/deck). The decks were randomized and ranked by 6 blinded investigators. GFP intensity was ranked from low (1) to high (11).

### LacZ staining

*OCR-Gal4* flies were crossed to *UAS-LacZ* female virgins. 5-6-day old progeny were collected and frozen at −80 °C. Frozen flies were allowed to equilibrate to -15-16 °C in a Leica CM1950 cryostat for at least 30 min. Then, whole flies were embedded in OCT cutting medium and sectioned at 30 μm. The sections were mounted on Superfrost Plus slides and incubated with 500 μL 2% X-Gal (20 mg/mL in DMF, Cat no. X1001-5, Zymo Research, Irvine, CA, USA) in 40 mL X-Gal staining solution (10 mM sodium phosphate buffer pH 7.2, 150 mM NaCl, 1 mM MgCl_2_, 10 mM K_4_[Fe^II^(CN)_6_], 10 mM K_3_[Fe^III^(CN)_6_], and 0.1% Triton X-100) overnight at 37 °C. The slides were washed 3X with 1X PBS and mounted with Prolong Diamond mounting medium (ThermoFisher). The labeled sections were imaged with an Olympus CKX53 inverted microscope. The final images were reconstructed in Adobe Photoshop v23.1.1.

### ATAC-seq library construction from OCR-Gal4 flies

ATAC-seq libraries were constructed from fly nuclei expressing *UAS-Stinger* driven by *3R579-Gal4* and *2L722-Gal4.* The nuclei were collected and the ATAC-seq libraries were constructed as described ^23, 45^. The libraries were sequenced with a Novaseq 6000 instrument (Illumina) using 50-bp paired-end reads and the resulting data were analyzed as described for the initial ATAC-seq libraries. OCRs with increased accessibility in *3R579-Gal4* and *2L722-Gal4* flies compared to *elav-* and *nSyb-Gal4* were selected by location in first introns, log_2_ fold change > 1, and adjusted *p* < 0.05. *OCR2*s were subcloned into pMDF as described above.

### Immunofluorescence

*OCR-Gal4* male flies were crossed to *UAS-myr::smGdP-V5* virgin females. Brains from 5-6 day old progeny were dissected in ice-cold PBS containing 0.5% Triton-X 100 (T-PBS) and placed in 4% paraformaldehyde in PBS (PFA) on ice. The brains were post-fixed for 20 min in fresh 4% PFA at room temperature on a nutator. After fixing, the tissue was washed 3X with 0.5% T-PBS (20 min each), blocked for at least 30 min with 5% normal goat serum in 0.5% T-PBS, and incubated at 4 °C for at least two nights with mouse anti-nc82 [1:20, Drosophila Studies Hybridoma Bank (DSHB) at the University of Iowa], AlexaFluor 647-conjugated mouse anti-V5 (clone SV5-Pk1, 1:300, BioRad), and rat anti-elav (clone 7E8A10, 1:700, DSHB) in 5% normal goat serum in 0.5% T-PBS. The brains were washed 3X with 0.5% T-PBS and incubated for at least two nights in goat anti-mouse AlexaFluor 488 (1:1000, ThermoFisher) and goat anti-rat AlexaFluor 594 (1:1000, ThermoFisher). After labeling, the brains were washed with 0.5% T-PBS for 20 min and post-fixed in 4% PFA in PBS for 4 h at room temperature on a nutator. The tissue was washed once in 0.5% T-PBS, 3X in 1X PBS (15 min each), and mounted on Superfrost Plus slides (VWR). The brains were dehydrated in 30, 50, 75, 95, and 3X 100% ethanol for 10 min each. The slides were cleared 3X in xylene (5 min each) and mounted with DPX (Sigma).

Male flies containing *OCR2*s (*OCR2-Flp;2L722-Gal4*) were crossed to *UAS-myr::GFP-FRT-STOP-FRT-smHA* female virgins for 24 h. The flies were flipped into a new vial, and were allowed to mate and lay eggs for 24 h. The parental flies were removed and the seeded vials were incubated at 25 °C for 24 hours to allow eggs to hatch and the larvae to develop to the first instar stage. The first-instar larvae were heat shocked at 37 °C for 1 h in a water bath. 24 h later, the larvae were heat-shocked again and allowed to recover for another 24 h. The larvae were heat-shocked a third time and allowed to develop normally afterward. Brains were dissected and stained as described above using mouse anti-nc82 (1:20), chicken anti-GFP (1:500, Cat. Millipore-Sigma), and AlexaFluor 647-conjugated mouse anti-HA (clone 6E2, 1:300, Cell Signaling Technologies, Danvers, MA, USA). Secondary antibodies were goat anti-mouse AlexaFluor 488 (1:1000, ThermoFisher) and goat anti-chicken AlexaFluor Plus 594 (1:1000, Cat. No. A32759, ThermoFisher). Immunostained brains were washed, post-fixed, and mounted as described above. All immunolabeled brains were imaged using the Leica SP8 white light laser confocal microscope at the University of Utah HSC Cell Imaging Core Facility. All images were processed using the Fiji distribution of ImageJ (v2.3.0/1.53q).^55^

### Sleep experiments

For sleep experiments, *OCR*-*Gal4* male flies were crossed to *UAS-TrpA1* female virgins. *OCR2-Flp;2L722-Gal4* male flies were crossed to *UAS-FRT-STOP-FRT-TrpA1* female virgin flies. The *OCR2*-expressing progeny were heat shocked to induce Flp activity as described above. Fly locomotor activity was monitored using the Drosophila Activity Monitor system (DAM3, Trikinetics, Waltham, MA). Flies were individually loaded into 65-mm glass tubes and placed within the DAM system. Flies were kept in an incubator with a 12-h light:dark cycle. Behavior was recorded for 2 days at 21 °C. Then, the temperature was shifted to 29 °C and activity was recorded for 2 more days. The sleep data was quantified using custom software ^56^ in MATLAB (MathWorks, Natick, MA). The data obtained for each sleep parameter at 21 °C was subtracted from the data from the same fly obtained at 29 °C before statistical analysis.

### Statistical analysis

Statistical analysis was performed in GraphPad 9 (version 9.4.1) or R (version 4.2.1) software. Annotated feature proportions were analyzed using Mann-Whitney U tests (whole flies vs. head neurons, Fig. 1B) or Kruskal-Wallis tests followed by Dunn’s multiple comparisons tests (head neurons vs. *2L722* or *3R579* neurons, Fig. 5C). GFP fluorescence intensity (Fig. 2) and reporter immunostaining (Fig. 3) were analyzed using Wilcoxon signed-rank tests. Differences in nighttime sleep were analyzed using Kruskal-Wallis tests followed by Dunn’s multiple comparisons test. Reporter labeling in Fig. 6 and Supplemental Fig. 4 was ranked by blinded investigators. The frequency of staining for each pattern was calculated and compared using Chi-squared tests. Differences were considered statistically significant at *p* < 0.05.

## SUPPLEMENTAL INFORMATION

**Supplemental Figure 1 O*C*R*-Gal4*-driven GFP fluorescence in whole flies.** Flies were imaged using the same microscope and camera settings. The values in white text are the randomized image numbers used for ranking.

**Supplemental Figure 2 LacZ staining in *OCR-Gal4;UAS-LacZ* flies.** Flies were imaged at 10X magnification. The scale bar shown for *2L009* indicates 500 μm and is the same for all images.

**Supplemental Figure 3 Neuron-derived *OCR-Gal4* flies drive reporter expression in neurons.** A-I) Expression of membrane-bound (myristoylated) V5 (green, encoded by *UAS-smV5*) was driven by each *OCR-Gal4*. Neuronal nuclei were stained for elav (magenta). V5 was stained in brains containing neuron-derived and whole fly-derived OCRs. Neuropil was stained with nc82. The OCR name is shown in the upper right corner of each image. Scale bar indicates 20 µm. The scale is the same for all images.

**Supplemental Figure 4 Representative OCR2 expression used for ranking.** A-J) Representative images of OCR2 expression in the brain. K) Dotplot showing the percentage of each pattern observed in each OCR2 fly line. Dot size indicates the number of observations of each pattern in each genotype, expressed as the total number of ranked brains (n = 60 for *2R236*, 75 for *2L945*, and 60 for control). Each image was independently ranked by 5 blinded researchers.

**Supplemental Figure 5 Differentially-accessible peaks detected by iterative ATAC-seq drive transgene expression in *3R579* neuron subtypes.** GFP expression (green) was driven by *3R579-Gal4*, while HA expression (magenta) was driven in neurons expressing both *3R579-Gal4* and *3R579*-derived *OCR2-FLPs*. The top panels show the anterior brain and the bottom panels show the posterior brain. Multiple images are shown for several *OCR2-Flp* lines due to incomplete *Flp* penetrance. Scale bar = 100 μm. All brains were imaged at the same magnification.

## Notes

### Competing Interest Statement

The authors have declared no competing interest.

## REFERENCES

1. Yuste, R. (2015). From the neuron doctrine to neural networks. Nat Rev Neurosci 16, 487–497. 10.1038/nrn3962.

2. Liu, C., Plaa_Jais, P.Y., Yamagata, N., Pfeiffer, B.D., Aso, Y., Friedrich, A.B., Siwanowicz, I., Rubin, G.M., Preat, T., and Tanimoto, H. (2012). A subset of dopamine neurons signals reward for odour memory in drosophila. Nature 488, 512–516. 10.1038/nature11304.

3. Aso, Y., Hattori, D., Yu, Y., Johnston, R.M., Iyer, N.A., Ngo, T.T.B., Dionne, H., Abbott, L.F., Axel, R., Tanimoto, H., and Rubin, G.M. (2014). The neuronal architecture of the mushroom body provides a logic for associative learning. eLife 3, e04577–e04577. 10.7554/ELIFE.04577.

4. Scheffer, L.K., Xu, C.S., Januszewski, M., Lu, Z., Takemura, S.-Y., Hayworth, K.J., Huang, G.B., Shinomiya, K., Maitlin-Shepard, J., Berg, S., et al. (2020). A connectome and analysis of the adult drosophila central brain. eLife 9. 10.7554/elife.57443.

5. Pfeiffer, B.D., Jenett, A., Hammonds, A.S., Ngo, T.T.B., Misra, S., Murphy, C., Scully, A., Carlson, J.W., Wan, K.H., Laverty, T.R., et al. (2008). Tools for neuroanatomy and neurogenetics in drosophila. Proceedings of the National Academy of Sciences of the United States of America 105, 9715–9720. 10.1073/pnas.0803697105.

6. Jenett, A., Rubin, G.M., Ngo, T.T.B., Shepherd, D., Murphy, C., Dionne, H., Pfeiffer, B.D., Cavallaro, A., Hall, D., Jeter, J., et al. (2012). A gal4-driver line resource for drosophila neurobiology. Cell Reports 2, 991–1001. 10.1016/j.celrep.2012.09.011.

7. Farkas, G., Leibovitch, B.A., and Elgin, S.C.R. (2000). Chromatin organization and transcriptional control of gene expression in drosophila. Gene 253, 117–136. 10.1016/S0378-1119(00)00240-7.

8. Marstrand, T.T., and Storey, J.D. (2014). Identifying and mapping cell-type-specific chromatin programming of gene expression. Proceedings of the National Academy of Sciences of the United States of America 111. 10.1073/PNAS.1312523111/SUPPL_FILE/SAPP.PDF.

9. Gui, Y., Grzyb, K., Thomas, M.H., Ohnmacht, J., Garcia, P., Buttini, M., Skupin, A., Sauter, T., and Sinkkonen, L. (2021). Single-nuclei chromatin profiling of ventral midbrain reveals cell identity transcription factors and cell-type-specific gene regulatory variation. Epigenetics and Chromatin 14, 1–20. 10.1186/S13072-021-00418-3/FIGURES/6.

10. Palmateer, C.M., Moseley, S.C., Ray, S., Brovero, S.G., and Arbeitman, M.N. (2021). Analysis of cell-type-specific chromatin modifications and gene expression in drosophila neurons that direct reproductive behavior. PLOS Genetics 17, e1009240–e1009240. 10.1371/JOURNAL.PGEN.1009240.

11. Rotem, A., Ram, O., Shoresh, N., Sperling, R.A., Goren, A., Weitz, D.A., and Bernstein, B.E. (2015). Single-cell chip-seq reveals cell subpopulations defined by chromatin state. Nat Biotechnol 33, 1165–1172. 10.1038/nbt.3383.

12. Winick-Ng, W., Kukalev, A., Harabula, I., Zea-Redondo, L., Szabo, D., Meijer, M., Serebreni, L., Zhang, Y., Bianco, S., Chiariello, A.M., et al. (2021). Cell-type specialization is encoded by specific chromatin topologies. Nature 599, 684–691. 10.1038/s41586-021-04081-2.

13. Buenrostro, J.D., Giresi, P.G., Zaba, L.C., Chang, H.Y., and Greenleaf, W.J. (2013). Transposition of native chromatin for fast and sensitive epigenomic profiling of open chromatin, DNA-binding proteins and nucleosome position. Nature Methods 10, 1213–1218. 10.1038/nmeth.2688.

14. Ackermann, A.M., Wang, Z., Schug, J., Naji, A., and Kaestner, K.H. (2016). Integration of atac-seq and rna-seq identifies human alpha cell and beta cell signature genes. Molecular Metabolism 5, 233–244. 10.1016/j.molmet.2016.01.002.

15. Grbesa, I., Tannenbaum, M., Sarusi-Portuguez, A., Schwartz, M., and Hakim, O. (2017). Mapping genome-wide accessible chromatin in primary human t lymphocytes by atac-seq. Journal of Visualized Experiments 2017, 56313–56313. 10.3791/56313.

16. Wang, J., Zibetti, C., Shang, P., Sripathi, S.R., Zhang, P., Cano, M., Hoang, T., Xia, S., Ji, H., Merbs, S.L., et al. (2018). Atac-seq analysis reveals a widespread decrease of chromatin accessibility in age-related macular degeneration. Nature Communications 9, 1–13. 10.1038/s41467-018-03856-y.

17. Bysani, M., Agren, R., Davegårdh, C., Volkov, P., Rönn, T., Unneberg, P., Bacos, K., and Ling, C. (2019). Atac-seq reveals alterations in open chromatin in pancreatic islets from subjects with type 2 diabetes. Scientific Reports 9. 10.1038/s41598-019-44076-8.

18. Fujiwara, S., Baek, S., Varticovski, L., Kim, S., and Hager, G.L. (2019). High quality atac-seq data recovered from cryopreserved breast cell lines and tissue. Scientific Reports 9, 1–11. 10.1038/s41598-018-36927-7.

19. Sinnamon, J.R., Torkenczy, K.A., Linhoff, M.W., Vitak, S.A., Mulqueen, R.M., Pliner, H.A., Trapnell, C., Steemers, F.J., Mandel, G., and Adey, A.C. (2019). The accessible chromatin landscape of the murine hippocampus at single-cell resolution. Genome Research 29, 857–869. 10.1101/gr.243725.118.

20. Aydin, B., Kakumanu, A., Rossillo, M., Moreno-Estellés, M., Garipler, G., Ringstad, N., Flames, N., Mahony, S., and Mazzoni, E.O. (2019). Proneural factors ascl1 and neurog2 contribute to neuronal subtype identities by establishing distinct chromatin landscapes. Nature Neuroscience 22, 897–908. 10.1038/s41593-019-0399-y.

21. Janssens, J., Aibar, S., Taskiran, II, Ismail, J.N., Gomez, A.E., Aughey, G., Spanier, K.I., De Rop, F.V., Gonzalez-Blas, C.B., Dionne, M., et al. (2022). Decoding gene regulation in the fly brain. Nature 601, 630–636. 10.1038/s41586-021-04262-z.

22. Brand, A.H., and Perrimon, N. (1993). Targeted gene expression as a means of altering cell fates and generating dominant phenotypes. Development 118, 401–415. 10.1242/DEV.118.2.401.

23. Merrill, C.B., Pabon, M.A., Montgomery, A.B., Rodan, A.R., and Rothenfluh, A. (2022). Optimized assay for transposase-accessible chromatin by sequencing (atac-seq) library preparation from adult drosophila melanogaster neurons. Sci Rep 12, 6043. 10.1038/s41598-022-09869-4.

24. Roy, S., Ernst, J., Kharchenko, P.V., Kheradpour, P., Negre, N., Eaton, M.L., Landolin, J.M., Bristow, C.A., Ma, L., Lin, M.F., et al. (2010). Identification of functional elements and regulatory circuits by drosophila modencode. Science 330, 1787–1797. 10.1126/science.1198374.

25. Gisselbrecht, S.S., Palagi, A., Kurland, J.V., Rogers, J.M., Ozadam, H., Zhan, Y., Dekker, J., and Bulyk, M.L. (2020). Transcriptional silencers in drosophila serve a dual role as transcriptional enhancers in alternate cellular contexts. Molecular Cell 77, 324–337.e328. 10.1016/j.molcel.2019.10.004.

26. Bischof, J., Maeda, R.K., Hediger, M., Karch, F., and Basler, K. (2007). An optimized transgenesis system for drosophila using germ-line-specific φc31 integrases. Proceedings of the National Academy of Sciences of the United States of America 104, 3312–3317. 10.1073/PNAS.0611511104/SUPPL_FILE/11511FIG4.JPG.

27. O’Brien, T., and Lis, J.T. (1993). Rapid changes in drosophila transcription after an instantaneous heat shock. Molecular and Cellular Biology 13, 3456–3463. doi:10.1128/mcb.13.6.3456-3463.1993.

28. Nern, A., Pfeiffer, B.D., and Rubin, G.M. (2015). Optimized tools for multicolor stochastic labeling reveal diverse stereotyped cell arrangements in the fly visual system. Proceedings of the National Academy of Sciences 112, E2967–E2976. 10.1073/PNAS.1506763112.

29. Hamada, F.N., Rosenzweig, M., Kang, K., Pulver, S.R., Ghezzi, A., Jegla, T.J., and Garrity, P.A. (2008). An internal thermal sensor controlling temperature preference in drosophila. Nature 454, 217–220. 10.1038/nature07001.

30. Donlea, J.M., Pimentel, D., and Miesenböck, G. (2014). Neuronal machinery of sleep homeostasis in drosophila. Neuron 81, 860–872. 10.1016/J.NEURON.2013.12.013.

31. Guo, F., Yu, J., Jung, H.J., Abruzzi, K.C., Luo, W., Griffith, L.C., and Rosbash, M. (2016). Circadian neuron feedback controls the drosophila sleep--activity profile. Nature 536, 292–297. 10.1038/nature19097.

32. Shalaby, N.A., Pinzon, J.H., Narayanan, A.S., Jin, E.J., Ritz, M.P., Dove, R.J., Wolfenberg, H., Rodan, A.R., Buszczak, M., and Rothenfluh, A. (2018). Jmjc domain proteins modulate circadian behaviors and sleep in drosophila. Sci Rep 8, 815. 10.1038/s41598-017-18989-1.

33. Shafer, O.T., and Keene, A.C. (2021). The regulation of drosophila sleep. Current Biology 31, R38–R49. 10.1016/j.cub.2020.10.082.

34. Fisher, Y.E., Yang, H.H., Isaacman-Beck, J., Xie, M., Gohl, D.M., and Clandinin, T.R. (2017). Flpstop, a tool for conditional gene control in drosophila. eLife 6. 10.7554/ELIFE.22279.

35. Duffy, J.B., Harrison, D.A., and Perrimon, N. (1998). Identifying loci required for follicular patterning using directed mosaics. Development 125, 2263–2271. 10.1242/DEV.125.12.2263.

36. Blair, S.S. (2003). Genetic mosaic techniques for studying drosophiladevelopment. Development 130, 5065–5072. 10.1242/DEV.00774.

37. Blankvoort, S., Witter, M.P., Noonan, J., Cotney, J., and Kentros, C. (2018). Marked diversity of unique cortical enhancers enables neuron-specific tools by enhancer-driven gene expression. Current Biology 28, 2103–2114.e2105. 10.1016/j.cub.2018.05.015.

38. Zhu, X., Ahmad, S.M., Aboukhalil, A., Busser, B.W., Kim, Y., Tansey, T.R., Haimovich, A., Jeffries, N., Bulyk, M.L., and Michelson, A.M. (2012). Differential regulation of mesodermal gene expression by drosophila cell type-specific forkhead transcription factors. Development 139, 1457–1466.

39. Bozek, M., Cortini, R., Storti, A.E., Unnerstall, U., Gaul, U., and Gompel, N. (2019). Atac-seq reveals regional differences in enhancer accessibility during the establishment of spatial coordinates in the drosophila blastoderm. Genome Research 29, 771–783. 10.1101/gr.242362.118.

40. Hrvatin, S., Tzeng, C.P., Nagy, M.A., Stroud, H., Koutsioumpa, C., Wilcox, O.F., Assad, E.G., Green, J., Harvey, C.D., Griffith, E.C., and Greenberg, M.E. (2019). A scalable platform for the development of cell-type-specific viral drivers. eLife 8. 10.7554/ELIFE.48089.

41. Graybuck, L.T., Daigle, T.L., Sedeño-Cortés, A.E., Walker, M., Kalmbach, B., Lenz, G.H., Morin, E., Nguyen, T.N., Garren, E., Bendrick, J.L., et al. (2021). Enhancer viruses for combinatorial cell-subclass-specific labeling. Neuron 109, 1449–1464.e1413. 10.1016/j.neuron.2021.03.011.

42. Luan, H., Diao, F., Scott, R.L., and White, B.H. (2020). The drosophila split gal4 system for neural circuit mapping. Front Neural Circuits 14, 603397. 10.3389/fncir.2020.603397.

43. Hsu, C.T., Choi, J.T.Y., and Sehgal, A. (2021). Manipulations of the olfactory circuit highlight the role of sensory stimulation in regulating sleep amount. Sleep 10.1093/sleep/zsaa265.

44. Cavey, M., Collins, B., Bertet, C., and Blau, J. (2016). Circadian rhythms in neuronal activity propagate through output circuits. Nature Neuroscience 19, 587–595. 10.1038/nn.4263.

45. Corces, M.R., Trevino, A.E., Hamilton, E.G., Greenside, P.G., Sinnott-Armstrong, N.A., Vesuna, S., Satpathy, A.T., Rubin, A.J., Montine, K.S., Wu, B., et al. (2017). An improved atac-seq protocol reduces background and enables interrogation of frozen tissues. Nature Methods 14, 959–962. 10.1038/nmeth.4396.

46. Martin, M. (2011). Cutadapt removes adapter sequences from high-throughput sequencing reads. EMBnet.journal 17, 10–10. 10.14806/ej.17.1.200.

47. Langmead, B., and Salzberg, S.L. (2012). Fast gapped-read alignment with bowtie 2. Nat Methods 9, 357–359. 10.1038/nmeth.1923.

48. Li, H., Handsaker, B., Wysoker, A., Fennell, T., Ruan, J., Homer, N., Marth, G., Abecasis, G., and Durbin, R. (2009). The sequence alignment/map format and samtools. Bioinformatics 25, 2078–2079.

49. Zhang, Y., Liu, T., Meyer, C.A., Eeckhoute, J., Johnson, D.S., Bernstein, B.E., Nussbaum, C., Myers, R.M., Brown, M., Li, W., and Liu, X.S. (2008). Model-based analysis of chip-seq (macs). Genome Biology 9, R137–R137. 10.1186/gb-2008-9-9-r137.

50. Ross-Innes, C.S., Stark, R., Teschendorff, A.E., Holmes, K.A., Ali, H.R., Dunning, M.J., Brown, G.D., Gojis, O., Ellis, I.O., Green, A.R., et al. (2012). Differential oestrogen receptor binding is associated with clinical outcome in breast cancer. Nature 481, 389–393. 10.1038/nature10730.

51. Love, M.I., Huber, W., and Anders, S. (2014). Moderated estimation of fold change and dispersion for rna-seq data with deseq2. Genome Biology 15, 1–21. 10.1186/S13059-014-0550-8/FIGURES/9.

52. Yu, G., Wang, L.-G., and He, Q.-Y. (2015). Chipseeker: An r/bioconductor package for chip peak annotation, comparison and visualization. Bioinformatics 31, 2382–2383. 10.1093/bioinformatics/btv145.

53. Ramirez, F., Ryan, D.P., Gruning, B., Bhardwaj, V., Kilpert, F., Richter, A.S., Heyne, S., Dundar, F., and Manke, T. (2016). Deeptools2: A next generation web server for deep-sequencing data analysis. Nucleic Acids Res 44, W160–165. 10.1093/nar/gkw257.

54. Robinson, J.T., Thorvaldsdóttir, H., Winckler, W., Guttman, M., Lander, E.S., Getz, G., and Mesirov, J.P. (2011). Integrative genomics viewer. Nature Biotechnology 29, 24–26. 10.1038/nbt.1754.

55. Schindelin, J., Arganda-Carreras, I., Frise, E., Kaynig, V., Longair, M., Pietzsch, T., Preibisch, S., Rueden, C., Saalfeld, S., Schmid, B., et al. (2012). Fiji: An open-source platform for biological-image analysis. Nature Methods 9, 676–682. 10.1038/nmeth.2019.

56. Vaccaro, A., Kaplan Dor, Y., Nambara, K., Pollina, E.A., Lin, C., Greenberg, M.E., and Rogulja, D. (2020). Sleep loss can cause death through accumulation of reactive oxygen species in the gut. Cell 181, 1307–1328.e1315. 10.1016/j.cell.2020.04.049.

