## Supplementary figures and images for "Iterative assay for transposase-accessible chromatin by sequencing to isolate functionally relevant neuronal subtypes"

### Supplemental Figures

Supplemental Fig. 1

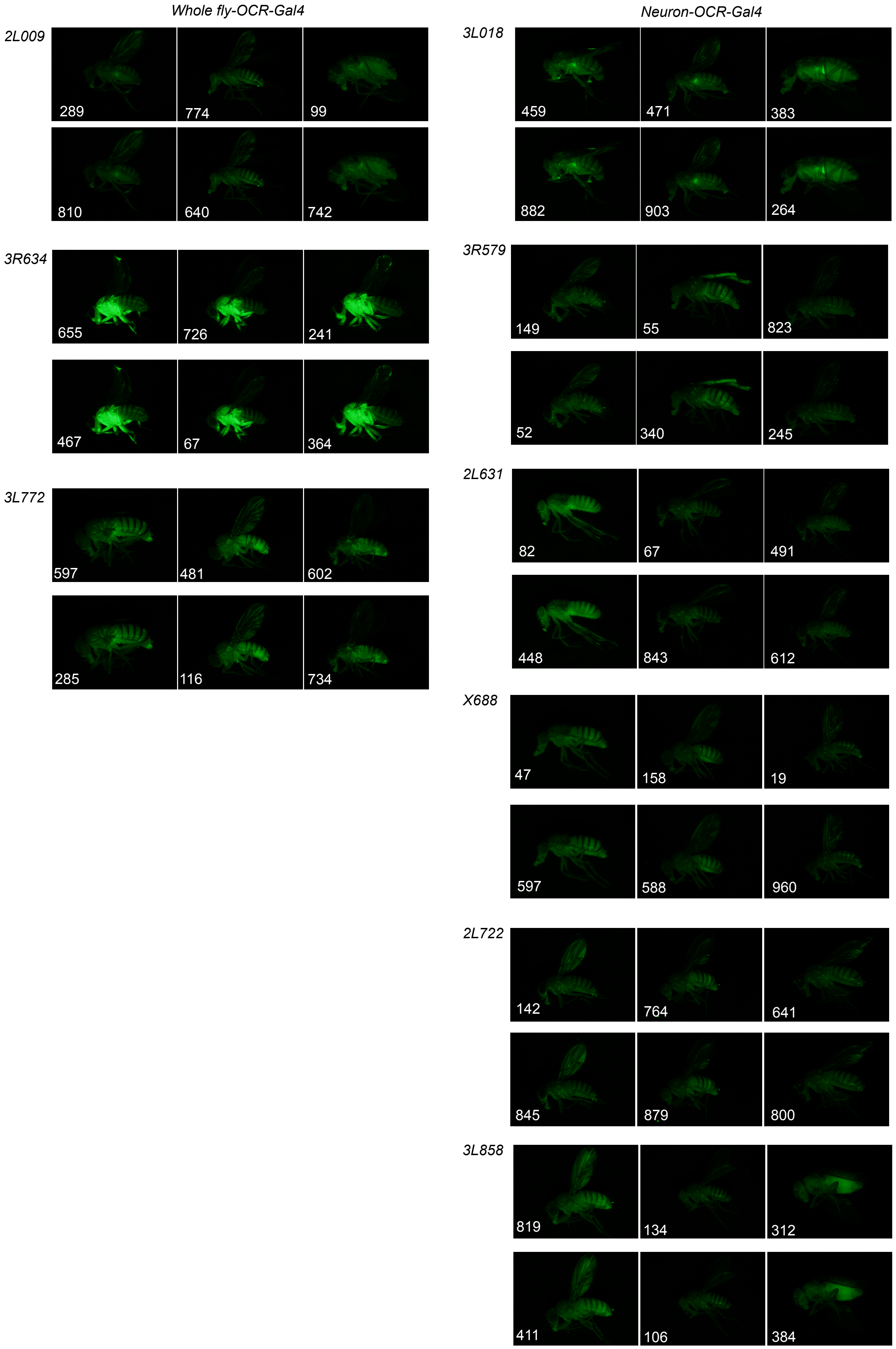

Supplemental Fig. 2

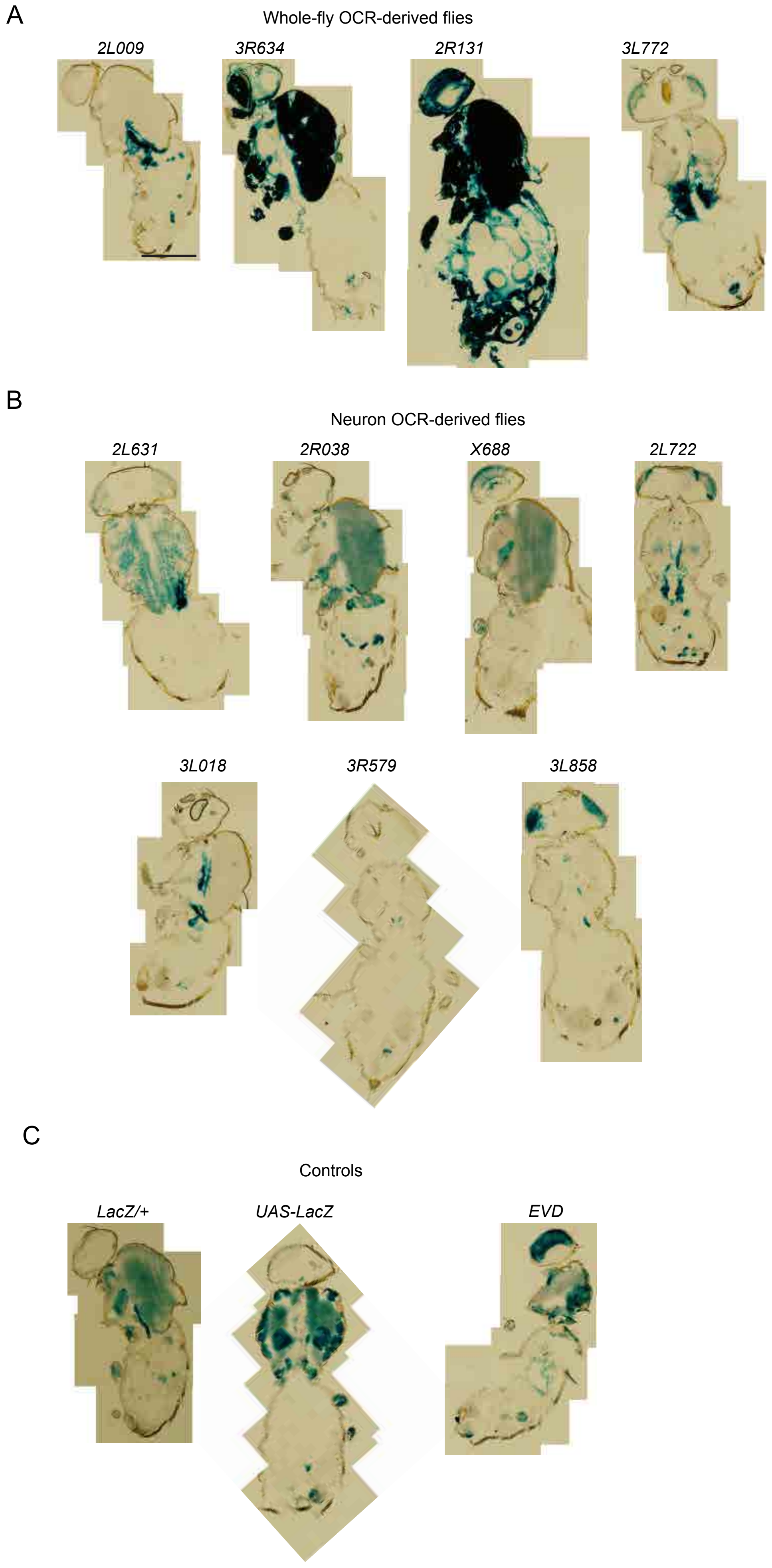

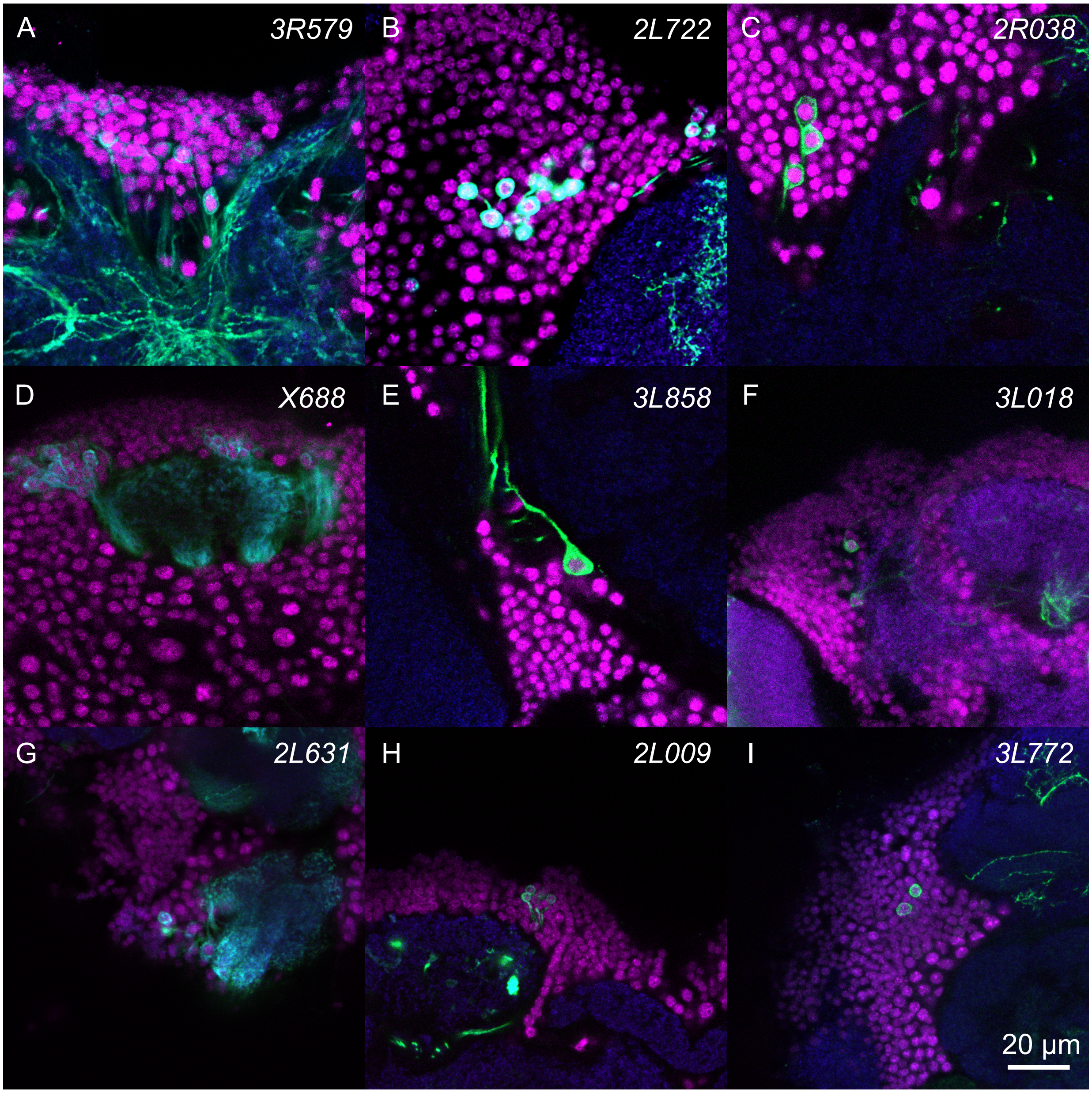

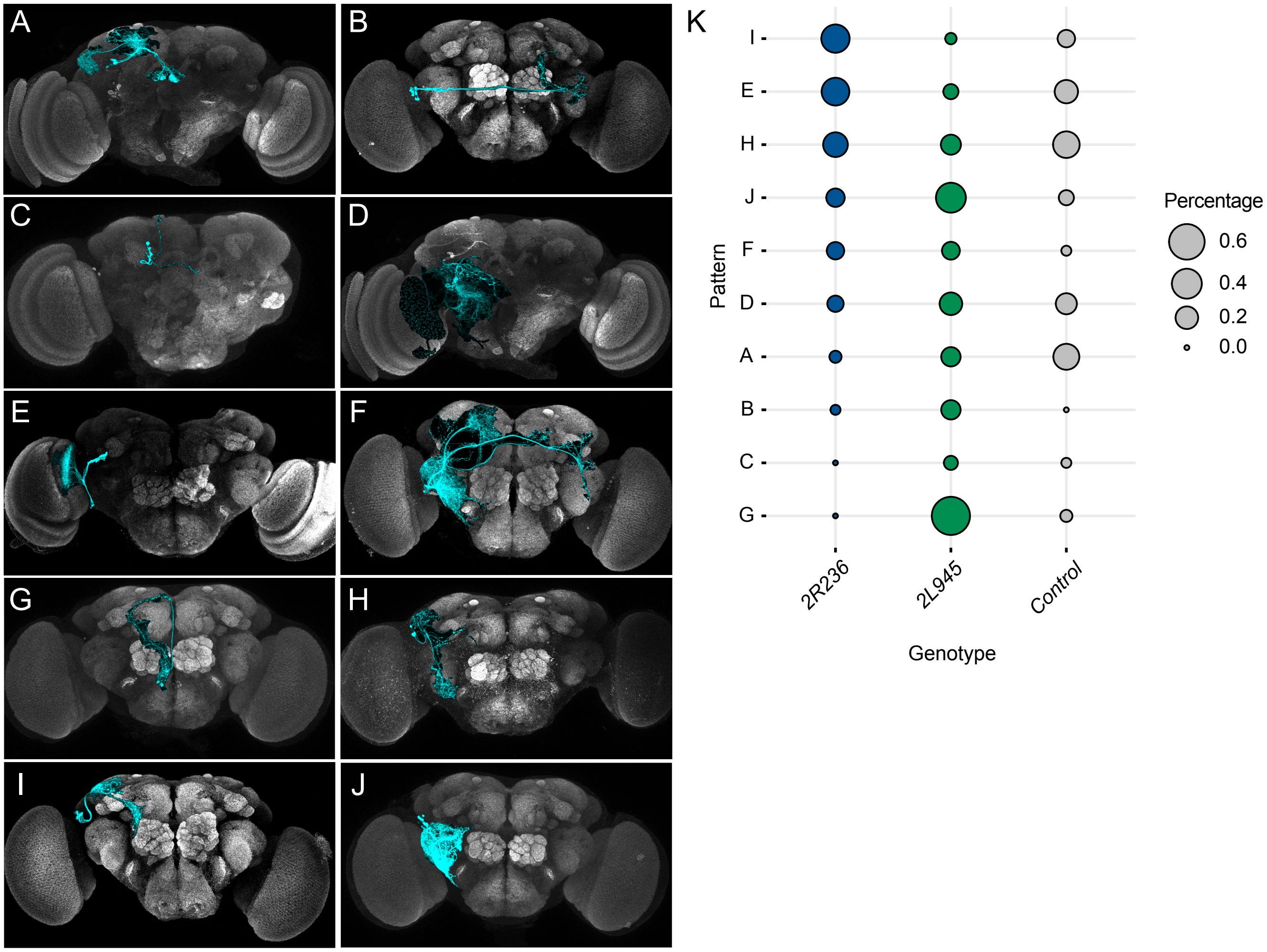

Supplemental Fig. 5

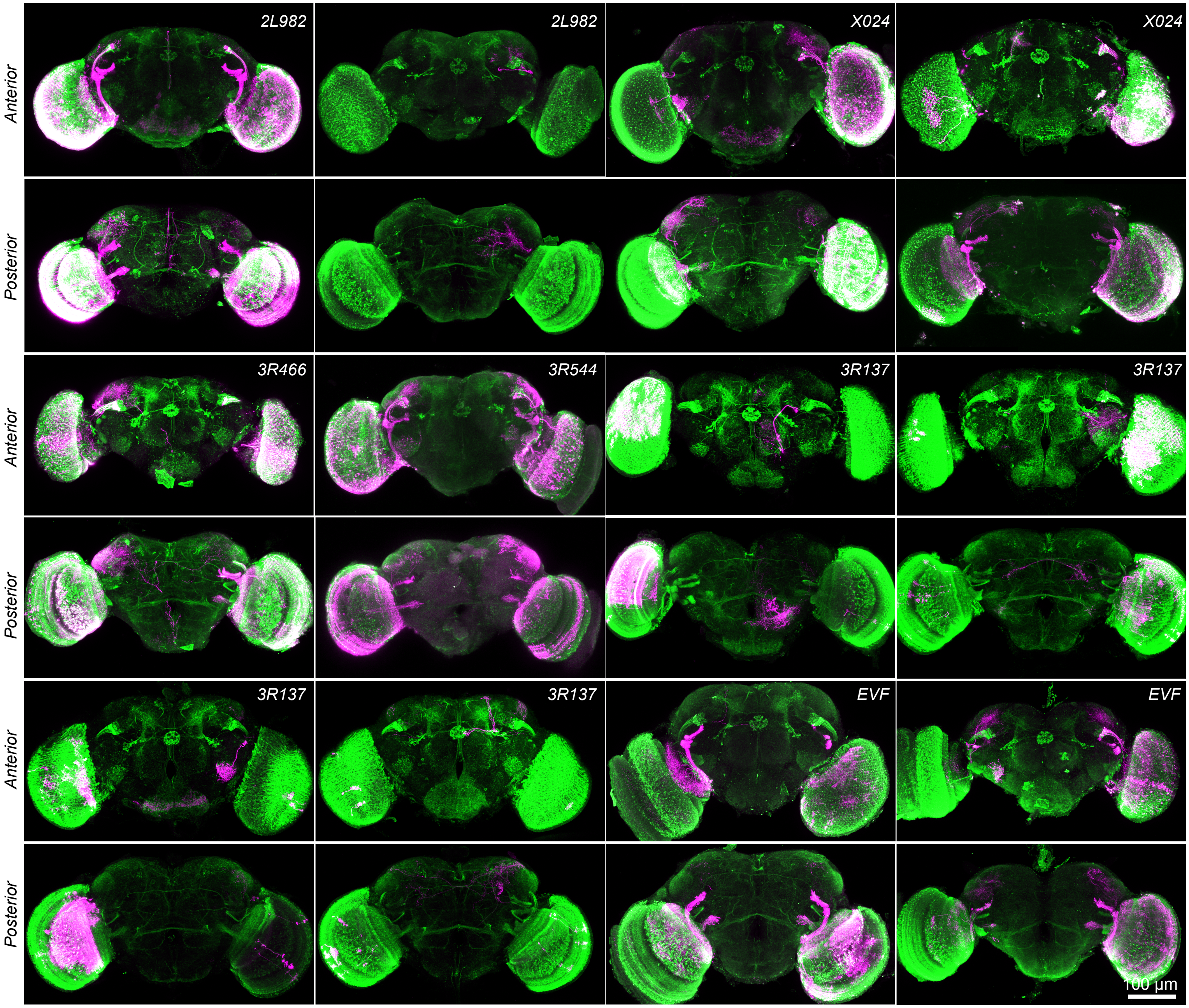
